# Broad cross-reactivity across sarbecoviruses exhibited by a subset of COVID-19 donor-derived neutralizing antibodies

**DOI:** 10.1101/2021.04.23.441195

**Authors:** Claudia A. Jette, Alexander A. Cohen, Priyanthi N.P. Gnanapragasam, Frauke Muecksch, Yu E. Lee, Kathryn E. Huey-Tubman, Fabian Schmidt, Theodora Hatziioannou, Paul D. Bieniasz, Michel C. Nussenzweig, Anthony P. West, Jennifer R. Keeffe, Pamela J. Bjorkman, Christopher O. Barnes

## Abstract

Many anti-SARS-CoV-2 neutralizing antibodies target the ACE2-binding site on viral spike receptor-binding domains (RBDs). The most potent antibodies recognize exposed variable epitopes, often rendering them ineffective against other sarbecoviruses and SARS-CoV-2 variants. Class 4 anti-RBD antibodies against a less-exposed, but more-conserved, cryptic epitope could recognize newly-emergent zoonotic sarbecoviruses and variants, but usually show only weak neutralization potencies. We characterized two class 4 anti-RBD antibodies derived from COVID-19 donors that exhibited broad recognition and potent neutralization of zoonotic coronavirus and SARS-CoV-2 variants. C118-RBD and C022-RBD structures revealed CDRH3 mainchain H-bond interactions that extended an RBD β-sheet, thus reducing sensitivity to RBD sidechain changes, and epitopes that extended from the cryptic epitope to occlude ACE2 binding. A C118-spike trimer structure revealed rotated RBDs to allow cryptic epitope access and the potential for intra-spike crosslinking to increase avidity. These studies facilitate vaccine design and illustrate potential advantages of class 4 RBD-binding antibody therapeutics.

## Introduction

The current SARS-CoV-2 pandemic is a crisis of immediate global concern, but two other zoonotic betacoronaviruses, SARS-CoV and MERS-CoV (Middle East Respiratory Syndrome), also resulted in epidemics within the last 20 years (de Wit et al., 2016). All three viruses likely originated in bats (Li et al., 2005; Zhou et al., 2021), with SARS-CoV and MERS-CoV having adapted to intermediary animal hosts, most likely palm civets (Song et al., 2005) and dromedary camels (Haagmans et al., 2014), respectively, prior to infection of humans. Serological surveys of people living near caves where bats carry diverse coronaviruses suggests direct transmission of SARS-CoV-like viruses (Wang et al., 2018), raising the possibility of future outbreaks resulting from human infection with SARS-like betacoronaviruses (sarbecoviruses).

Coronaviruses encode a trimeric spike glycoprotein (S) that serves as the machinery for fusing the viral and host cell membranes (Fung and Liu, 2019). The first step in fusion is contact of S with a host receptor. The receptor-binding domains (RBDs) at the apex of the S trimers of SARS-CoV-2, SARS-CoV, HCoV-NL63, and some animal coronaviruses utilize angiotensin-converting enzyme 2 (ACE2) as their receptor (Hoffmann et al., 2020; Li et al., 2003; Zhou et al., 2020b). RBDs can adopt either ‘down’ or ‘up’ conformations, with ACE2 binding only possible to RBDs in an ‘up’ conformation (Kirchdoerfer et al., 2016; Li et al., 2019; Walls et al., 2020; Walls et al., 2016; Wrapp et al., 2020; Yuan et al., 2017). A phylogenetic tree of the relationship between coronavirus S protein RBDs shows that sarbecovirus RBDs form a separate branch (Figure 1A).

**Figure 1.**
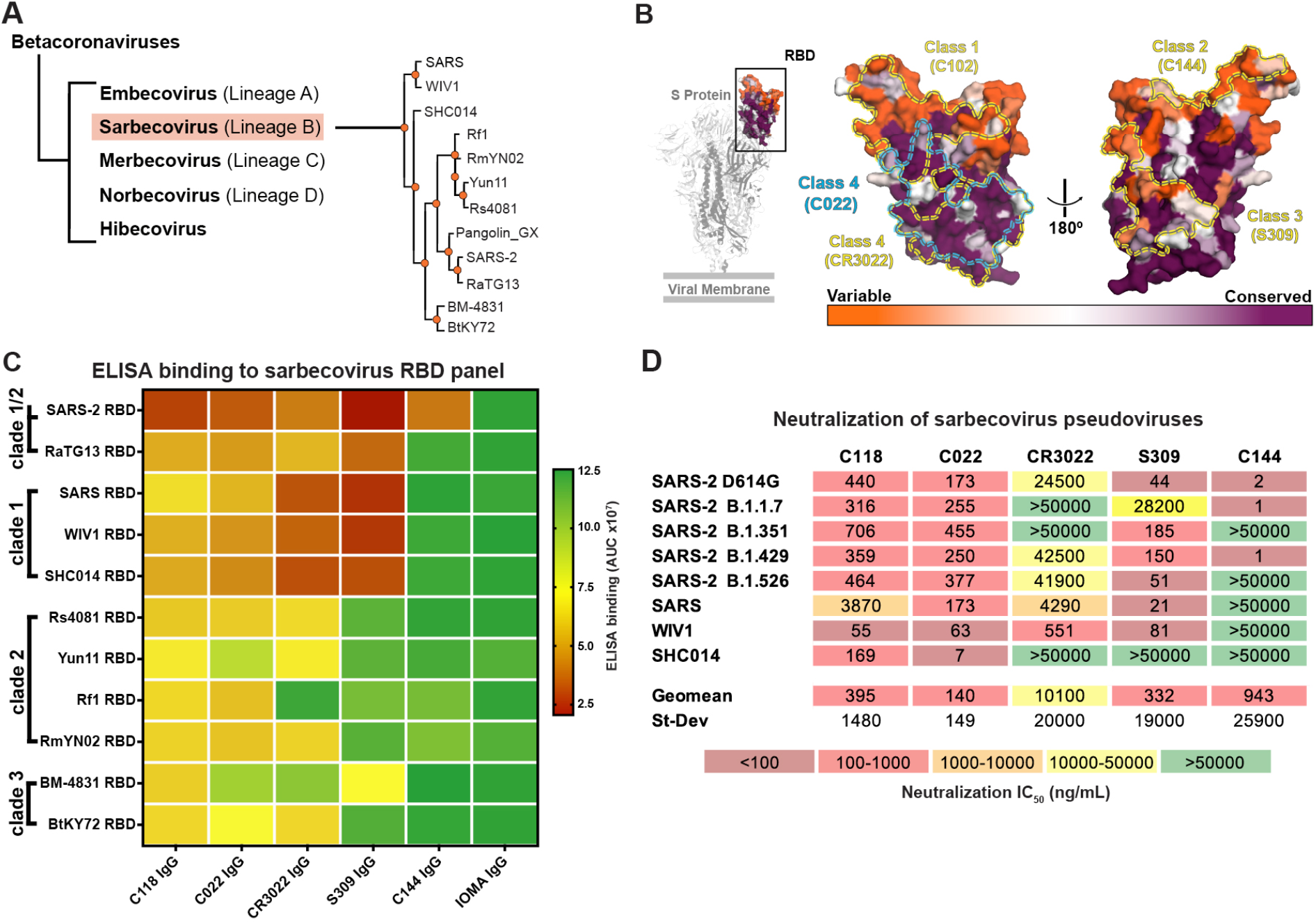
C118 and C022 show diverse binding and neutralization of sarbecoviruses. (A) Sarbecovirus (Lineage B) phylogenetic tree classified based on RBD sequence conservation. (B) Left: Cartoon rendering of SARS-CoV-2 S trimer (PDB 6VYB) showing location of ‘up’ RBD (surface, orange and purple). Right: Amino acid sequence conservation of 12 RBDs calculated as described (Landau et al., 2005) plotted on a surface representation of a SARS-CoV-2 RBD structure (PDB 7BZ5). Primary RBD epitopes for the indicated representatives from defined classes of RBD-binding antibodies (class 1-4) (Barnes et al., 2020a) are indicated as yellow dotted lines (PDB 7K90, 6W41, 7JX3,7K8M). C022 epitope indicated as blue dotted line. (C) Comparison of binding of the indicated monoclonal IgGs to a panel of sarbecovirus RBDs from ELISA data shown as area under the curve (AUC) values. Data presented are mean AUC values from two independent experiments. IOMA IgG is an anti-HIV-1 antibody serving as a negative control (Gristick et al., 2016). (D) Neutralization IC_50_ values for the indicated IgGs against SARS-CoV-2 (D614G version of the original variant (GenBank: NC_045512)), SARS-CoV-2 variants of concern, and other ACE2-tropic sarbecovirus pseudoviruses. Geomean = geometric mean IC_50_ in which IC_50_ values >50000ng/mL were entered as 50000 ng/mL for the calculation. SD = standard deviation. IC_50_ values are means of 2-7 independent experiments.

Consistent with their obligate role in viral entry, sarbecovirus S trimers are the primary targets of neutralizing antibodies (Brouwer et al., 2020; Cao et al., 2020; Fung and Liu, 2019; Kreer et al., 2020; Liu et al., 2020b; Robbiani et al., 2020; Rogers et al., 2020; Seydoux et al., 2020; Shi et al., 2020; Zost et al., 2020b), with many focusing on the RBD (Barnes et al., 2020a; Barnes et al., 2020b; Brouwer et al., 2020; Cao et al., 2020; Kreer et al., 2020; Liu et al., 2020b; Pinto et al., 2020; Robbiani et al., 2020; Rogers et al., 2020; Seydoux et al., 2020; Zost et al., 2020a). Structural analysis of the binding epitopes of anti-SARS-CoV-2 RBD antibodies enabled their classification into four initial categories: class 1, derived from *VH3-53*/*VH3-63* germlines and including a short heavy chain complementarity determining region 3 (CDRH3) that bind an epitope overlapping with the ACE2 binding site and only recognize ‘up’ RBDs; class 2, whose epitope also overlaps with the ACE2 binding site, but which can bind to both ‘up’ and ‘down’ RBD conformations; class 3, which bind to the opposite side of ‘up’ and ‘down’ RBDs adjacent to an N-glycan attached to residue N343; and class 4, which are often weakly neutralizing antibodies that target a cryptic epitope facing the interior of the spike protein on ‘up’ RBDs (Barnes et al., 2020a) (Figure S1).

Potent anti-SARS-CoV-2 neutralizing antibodies are typically class 1 or class 2 anti-RBD antibodies that block the ACE2 binding site (Barnes et al., 2020a; Dejnirattisai et al., 2021; Lee et al., 2021; Liu et al., 2020b; Piccoli et al., 2020a; Tortorici, 2020). Since class 1 and class 2 RBD epitopes are not well conserved (Figure 1B), antibodies in these classes are unlikely to strongly cross-react across sarbecovirus RBDs. However, an in vitro-selected variant of an ACE2 blocking antibody isolated from a SARS-infected survivor exhibited increased cross-reactive properties, showing neutralization of SARS-CoV-2 and other betacoronaviruses (Rappazzo et al., 2021). In general, however, as isolated from infected donors, class 3 and class 4 RBD-binding antibodies are better prospects for neutralizing across multiple strains and thereby potentially protecting against emergent sarbecoviruses. Indeed, S309, a class 3 anti-RBD antibody isolated from a SARS-CoV–infected donor, demonstrated cross-reactive neutralization of SARS-CoV-2 (Pinto et al., 2020). Furthermore, reports of class 4 human antibodies that exhibit cross-reactive binding and neutralization amongst sarbecoviruses (Liu et al., 2020a; Starr et al., 2021a; Tortorici et al., 2021) suggest that further investigation of antibodies from COVID-19 convalescent donors could lead to discoveries of potent and broadly cross-reactive class 4 antibodies that recognize the highly-conserved, ‘cryptic’ RBD epitope.

Here we investigated C118 and C022, two class 4 human antibodies isolated from COVID-19 donors (Robbiani et al., 2020) that show breadth of binding and neutralization across sarbecoviruses and SARS-CoV-2 variants of concern. We report crystal structures of C118 complexed with SARS RBD and C022 complexed with SARS-CoV-2 RBD, which revealed interactions with a conserved portion of the RBD in common with interactions of previously-described cross-reactive but more weakly-neutralizing class 4 antibodies; e.g., CR3022 (Huo et al., 2020; Yuan et al., 2020a; Yuan et al., 2020b), S304/S2A4 (Piccoli et al., 2020, Cell), and EY6A (Zhou et al., 2020a). Unlike these class 4 anti-RBD antibodies, C118 and C022 also occlude portions of the ACE2 binding site to facilitate more potent neutralization. A single-particle cryo-EM structure of a C118-S trimer complex demonstrated binding of C118 to an intact trimer, revealing an S configuration with increased separation between the RBDs than found in class 1-3 Fab-S or ACE2-S trimer structures, and revealed the potential for intra-spike crosslinking. These results define a cross-reactive class 4-like epitope on sarbecovirus RBDs that can be targeted in vaccine design and illustrate a mechanism by which the cryptic RBD epitope can be accessed on intact CoV S trimers.

## Results

### C022 and C118 IgGs recognize and neutralize diverse sarbecoviruses, including SARS-CoV-2 variants

From a survey to identify cross-reactive monoclonal antibodies isolated from SARS-CoV-2– infected donors from the New York area (Robbiani et al., 2020), we found antibodies isolated from different donors, C118 (*VH3-30/VL4-69*-encoded) and C022 (*VH4-39/VK1-5-*encoded), that recognized a diverse panel of 12 sarbecovirus RBDs spanning clades 1, 1/2, 2 and 3 (Figure 1). As evaluated by enzyme-linked immunosorbent assay (ELISA), C118 bound to RBDs from all sarbecoviruses tested, and C022 bound to all but two RBDs, similar to the class 4 anti-RBD antibody CR3022 (Figure 1C). By comparison, the cross-reactive class 3 anti-SARS RBD antibody S309 (Pinto et al., 2020) recognized half of the set of sarbecovirus RBDs, and C144, a more potent SARS-CoV-2 class 2 neutralizing antibody (Robbiani et al., 2020), bound to the SARS-CoV-2 RBD but not to RBDs from the other 11 sarbecovirus strains (Figure 1C).

To further define the C022 and C118 antibody epitopes, we evaluated binding of C118 and C022 to a panel of RBDs with mutations chosen from circulating variants that conferred resistance to one or more classes of anti-RBD antibodies (Li et al., 2020; Starr et al., 2021b; Weisblum et al., 2020). We also assessed binding to RBD substitutions identified in the B.1.1.7 and B.1.351 SARS-CoV-2 variants of concern (Rambaut et al., 2020; Tegally et al., 2020), and to mutations in the MA10 mouse-adapted SARS-CoV-2 virus (Leist et al., 2020). Relative to wild-type RBD, C118, C022, CR3022 and S309 demonstrated a similar binding profile with respect to the RBD substitutions tested and exhibited a broader range of binding to the RBD mutants than did the more potent class 2 C144 antibody (Figure 1C and Figure S2A). Collectively, the ELISA binding data suggested that C022 and C118 recognize a highly-conserved epitope and are therefore likely to be class 4 anti-RBD antibodies.

We next measured neutralization potencies using an in vitro pseudovirus-based assay that quantitatively correlates with authentic virus neutralization (Schmidt et al., 2020) to evaluate SARS-CoV-2, SARS-CoV-2 RBD mutants, SARS-CoV-2 variants (Annavajhala et al., 2021; Faria et al., 2021; Rambaut et al., 2020; Tegally et al., 2020; Voloch et al., 2020; West et al., 2021; Zhang et al., 2021), and sarbecovirus strains known to infect human ACE2-expressing target cells (SARS-CoV-2, SARS-CoV, WIV1, SHC104, WIV16, Pangolin GD and Pangolin GX) (Figure 1D and Figure S2B-D). Against a panel of SARS-CoV-2 pseudotyped viruses harboring single amino acid RBD substitutions, C118 and C022 neutralized all viruses with potencies similar to ‘wt’ SARS-CoV-2, consistent with the results obtained in ELISA binding assays (S gene with D614 residue; GenBank: NC_045512) (Figure S2). For comparisons with SARS-CoV-2 variants of concern, the S gene we used to make ‘wt’ SARS-CoV-2 pseudovirus included the D614G substitution in the context of the Wuhan-Hu-1 spike (Korber et al., 2020), resulting in a 2-4–fold reduction in IC_50_s for C022 and C118 antibodies (Figure 1D).

We found that C118 and C022 IgGs neutralized all four SARS-CoV-2 variants and all ACE2-tropic sarbecoviruses with 50% inhibitory concentrations (IC_50_ values) of <1 µg/mL, with the exception of C118, which inhibited SARS-CoV-pseudotyped viruses less efficiently (IC_50_ = ∼4.5 µg/mL) (Figure 1D and Figure S2B-D). By contrast, the class 4 anti-RBD antibody CR3022 showed weak or no neutralization against the majority of pseudoviruses tested, with the exception of SARS-CoV (IC_50_ ∼1.1 µg/mL) and WIV1 (IC_50_ ∼0.6 µg/mL). The class 3 S309 antibody showed strong neutralization potencies (IC_50_s between 16 ng/mL and 120 ng/mL) against all viruses with the exceptions of the B.1.1.7 SARS-CoV-2 variant of concern and SHC014. The class 2 anti-RBD antibody C144 was highly potent against SARS-CoV-2 and the B.1.1.7 and B.1.429 variants (IC_50_s between 1 ng/mL and 2 ng/mL), but did not neutralize the other SARS-CoV-2 variants or sarbecoviruses. Taken together, of the IgGs evaluated, C118 and C022 exhibited the greatest breadth of sarbecovirus neutralization (Figure 1D and Figure S2), consistent with their broad cross-reactive binding profile demonstrated by ELISA (Figure 1C and Figure S2A).

### Crystal structures of C022-RBD and C118-RBD reveal class 4 RBD interactions and conservation of epitope residues

To understand the mechanism underlying the breadth of neutralization of C022 and C118, we solved structures of complexes between C118 Fab bound to SARS-CoV RBD and C022 bound to SARS-CoV-2 RBD to resolutions of 2.7Å and 3.2Å, respectively, chosen based on which complexes formed well-ordered crystals (Figure 2A,B and Table S1).

**Figure 2.**
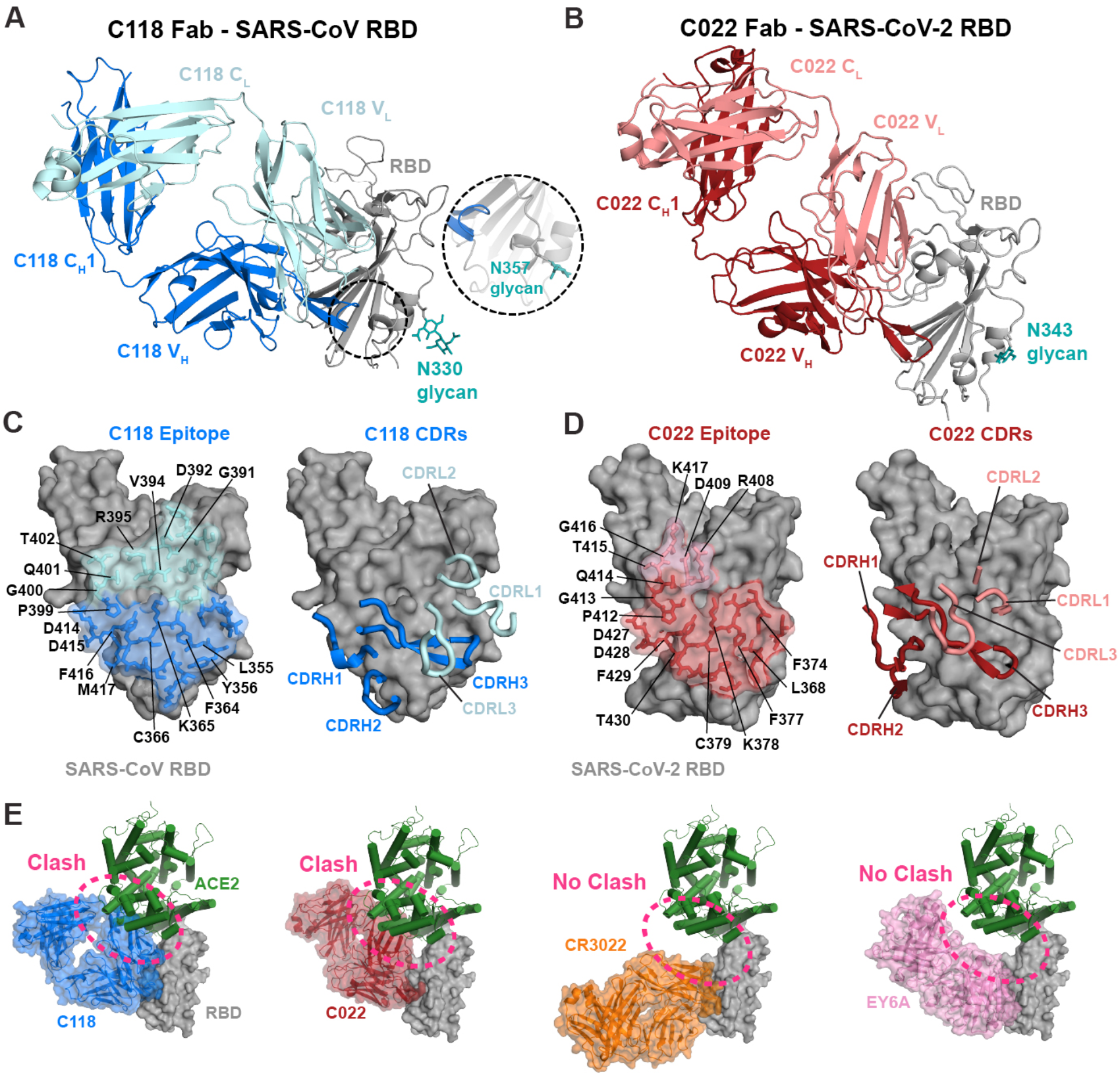
Crystal structures of C022 and C118 Fabs bound to RBDs reveal class 4-like RBD binding. (A,B) Cartoon renderings of crystal structures of (A) C0118 Fab complexed with SARS-CoV RBD, and (B) C022 Fab complexed with SARS-CoV-2 RBD. Dashed circle shows location of SARS-CoV N357_RBD_ residue, with the inset showing the N357_RBD_ asparagine and glycan modeled based on the SARS-CoV spike-S230 structure (PDB 6NB6). (C,D) CDR loops and RBD epitope residues of (C) C118 Fab and (D) C022 Fab overlaid on RBDs represented as gray surfaces with stick representations of epitope residues. Framework region residues, which account for some of the contacts for both antibodies, are not shown in right panels. (E) Comparison of Fab poses for binding to an RBD-ACE2 complex. C118 Fab (blue), C022 Fab (red), CR3022 Fab (PDB 6W41; orange), and EY6A Fab (PDB 6CZC pink) modeled onto an ACE2-RBD structure (PDB 6M0J; RBD shown as a gray surface and ACE2 shown as a green cartoon).

The C118-RBD and C022-RBD structures showed that both Fabs recognize an epitope that is highly-conserved among sarbecoviruses at the base of the RBD (Figure 1B), which is exposed only in ‘up’ RBD conformations as first described for the class 4 RBD-binding antibodies CR3022 (Huo et al., 2020; Yuan et al., 2020a; Yuan et al., 2020b) and EY6A (Zhou et al., 2020a). C022 and C118 use four of six complementarity-determining region (CDR) loops to interact with an epitope that extends towards the RBD ridge near the ACE2 binding site, and in the case of C022, includes an overlapping interacting residue (K417_RBD_) (Figure 2C,D). In both structures, CDRH3 loops, CDRL2 loops, and portions of FWRL3 mediate the majority of RBD contacts and establish extensive polar and van der Waals interactions with RBD residues (Figure 2C,D), accounting for 71% of epitope buried surface area (BSA) on the RBD for the C022-RBD and C118-RBD structures, respectively (Table S2). No contacts were made in either complex with the N343_RBD_ *N*-glycan (SARS-CoV-2 S numbering). SARS-CoV contains an additional potential *N*-linked glycosylation site at N357_RBD_ (SARS-CoV S numbering), which if glycosylated, would not be contacted by C118, a favorable feature for cross-reactive recognition given that this potential *N*-linked glycosylation site is conserved in all S protein sequences except for SARS-CoV-2 (Figure 2A).

Overlaying the RBDs of our Fab-RBD structures with the RBD of the ACE2-RBD structure (PDB 6M0J) showed that the binding poses of both C118 and C022 placed the V_L_ domain of each Fab in a position that would clash with concurrent ACE2 binding, in contrast to the CR3022 and EY6A binding poses (Figure 2E). This binding orientation would sterically prevent RBD-ACE2 interactions, as has been suggested for other class 4 anti-RBD antibodies (Liu et al., 2020a; Piccoli et al., 2020b). To verify direct competition with ACE2, we conducted a competition experiment using surface plasmon resonance (SPR). SARS-CoV-2 RBD was coupled to a biosensor chip, an RBD-binding IgG was injected, and then soluble ACE2 was injected over the RBD-IgG complex. C118, C022, and C144 IgGs all inhibited binding of ACE2, which did not bind to the IgG-RBD complex. In contrast, ACE2 bound the CR3022-RBD complex (Figure S3). These results are consistent with competition for C118, C022, and C144 for ACE2 binding to RBD, but no competition for CR3022, suggesting a primary neutralization mechanism for C022 and C118 that prevents spike attachment to host cell ACE2 receptors.

### Features of C118 and C022 recognition of the class 4 epitope

Class 4 RBD-binding antibodies contact a common epitope at the base of the RBD that is distant from the ACE2-binding site (Figure 3A). The epitopes of three class 4 antibodies, C118, C022, and COVA1-16, also includes a patch reaching towards the ridge on the left side of the RBD as depicted in Figure 3A.

**Figure 3.**
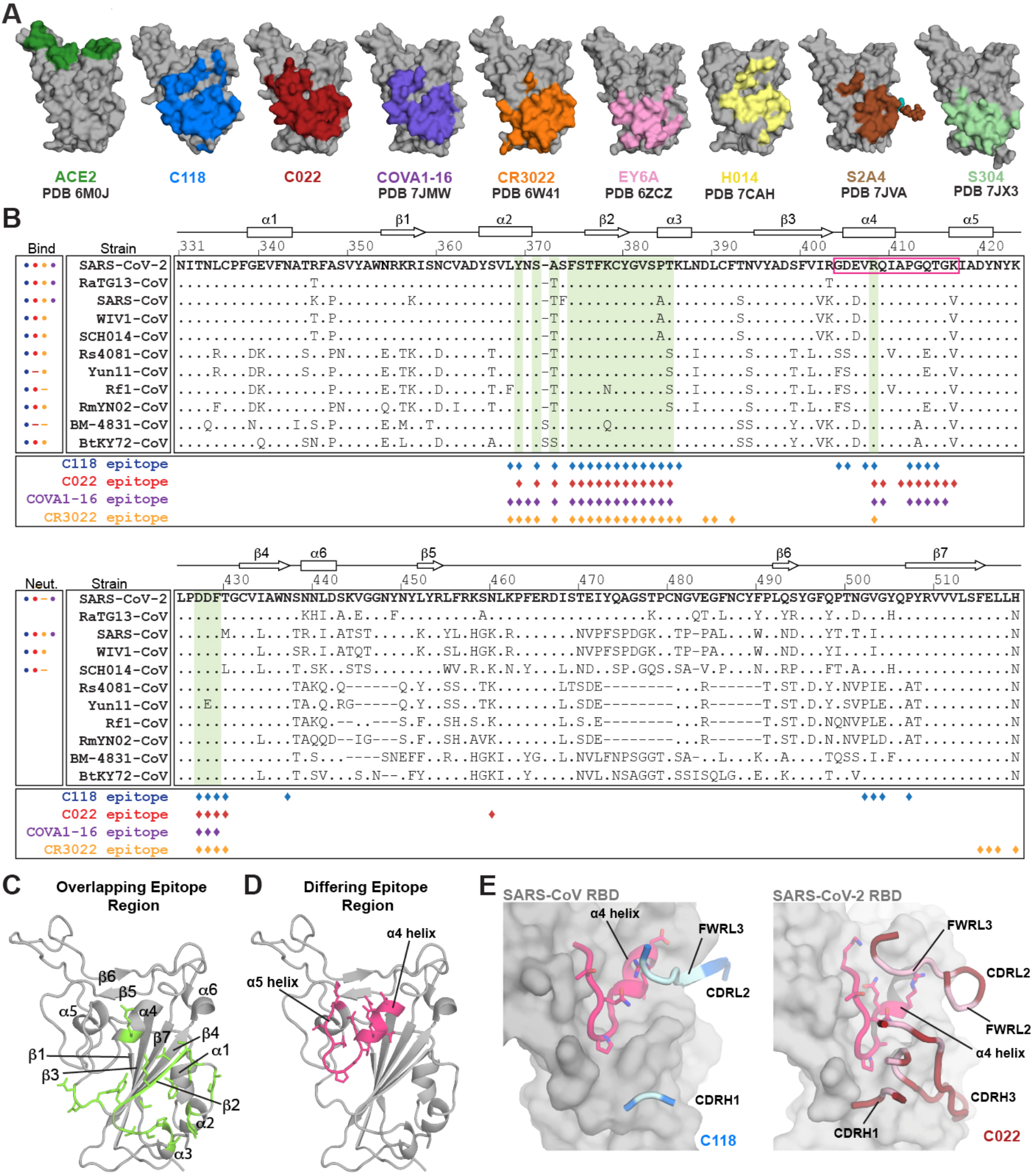
The C118 and C022 epitopes include a conserved RBD helix. (A) Epitopes for ACE2 and monoclonal antibodies calculated from analyses of structures of RBD or S trimer complexes (human antibodies isolated from COVID-19 patients are C118, C022, COVA1-16, EY6A, and S2A4). RBDs shown are derived from SARS-CoV-2 except for the C118 panel, which is SARS-CoV RBD. (B) Alignment of sequences for sarbecovirus RBDs (residue numbering for SARS-CoV-2 RBD). Secondary structure for SARS-CoV-2 RBD shown above alignment. Dots designate binding or neutralization for C118 (blue), C022 (red), or CR3022 (orange) for each strain. Diamonds designate RBD epitope residues for C118 binding to SARS-CoV (blue) and C022 (red) or CR3022 (orange) binding to SARS-CoV-2. Left boxes show binding by ELISA or neutralization of pseudovirus for each antibody for each strain; data for COVA1-16 from (Liu et al., 2020a). Circles show binding or neutralization, blank spaces designate not tested, and dashes designate no binding or neutralization. Shadings in the sequence alignment indicate conserved portions of epitope (green). Colored boxes show differing portion of epitope covering the α4 helix and following loop (pink. (C) Cartoon representation of SARS-CoV-2 RBD (gray) showing overlapping antibody-interacting residues (green) as sticks in epitopes for C118, C022, COVA1-16, and CR3022 (corresponding to green shading in panel B). (D) Cartoon representation of SARS-CoV-2 RBD (gray) showing α4 helix and following (sticks, pink) that differ in their contacts with C118, C022, COVA1-16, and CR3022 (pink shading in panel B). (E) Cartoon representation of RBDs showing α4 region of RBD and C118 (left) or C022 (right) interacting loops with interacting Fab residues in light blue (C118) and light pink (C022).

To compare the C118 and C022 epitopes with epitopes of other class 4 anti-RBD antibodies, we analyzed RBD residues contacted by C118, C022, COVA1-16, and CR3022 on aligned sequences of sarbecovirus RBDs (Figure 3B). Sequence conservation among sarbecoviruses at the C022 and C118 epitopes involves a majority of residues that are strictly-conserved or conservatively-substituted between SARS-CoV-2 and other RBDs (Figure 3B), likely explaining the broad cross-reactivity observed for these antibodies (Figure 1C). Comparison of the C118 and C022 epitopes showed a majority of recognized RBD residues are shared between the two antibodies (70% of C118 epitope also contacted by C022) (Figure 3B). CR3022 contacted a similar number of residues as C118 and C022, including the conserved patch at the RBD base (Figure 3B,C); however, a region from 404-417_RBD_ that comprises an unstructured loop and the α4 helix above an internal RBD β-sheet contained only a single CR3022 contact residue (R408_RBD_) and was not contacted by antibodies EY6A, S2A4 and S304; whereas C118, C022, and COVA1-16 showed contacts with this region (Figure 3B,D).

The α4 helix is proximal to the ACE2 receptor-binding motif and has less sequence conservation across the 12 sarbecoviruses (Figure 3B). To accommodate binding in this region, C118 uses insertions in its FWRL3 (54B-56_LC_) to form a β-strand adjacent to the α4 helix, establishing both side chain and backbone interactions (Figure 3E – left panel). C022 showed similar binding in this region but used non-contiguous CDRH1, CDRH3, and CDRL2 loops (Figure 3E – right panel). C022 contacts were located more to the C-terminal end of the α4 helix than the C118 contacts and encompassed the disordered RBD loop that includes the ACE2-interacting residue K417_RBD_ (K404_RBD_ in SARS-CoV) (Lan et al., 2020) (Figure 3E – right panel). Additioonally, C022 buried more surface area on RBD in this region than C118 (323Å^2^ vs 150Å^2^). Four of eight and five of nine RBD contacts for C118 and C022, respectively, were fully conserved among sarbecoviruses (Figure 3B), suggesting that interactions in this region may be possible with other sarbecoviruses. In particular, the conserved residue R408_RBD_ (R395_RBD_ in SARS-CoV) was contacted by both antibodies and alone was responsible for 94Å^2^ and 95Å^2^ of BSA buried on the RBDs for C118 and C022, respectively. Despite both C118 and C022 engaging the α4 helix and residue R408_RBD_, mutations at this position known to affect class 1 and class 4 anti-RBD antibodies (Greaney et al., 2021) had no effect on these antibodies (Figure S2A). Overall, engagement of the α4 helix region provided 16% (C118) and 36% (C022) of the BSA buried on RBD, and extended their epitopes past the cryptic epitope to bind adjacent to or overlapping with the ACE2 binding site.

### Shared features of the C022 and COVA1-16 class 4 anti-RBD antibodies

The C022 epitope on RBD closely resembles the epitope of COVA1-16 (Figure 3A,B), a class 4 antibody isolated from a SARS-CoV-2 convalescent donor derived from *VH1-46/VK1-33* V-gene segments (Brouwer et al., 2020) (Figure S4). Yet, COVA1-16 weakly neutralized (>1µg/mL) SARS-CoV and SARS-CoV-2 variants (Liu et al., 2021), in contrast with C022 neutralization (Figure 1D). After superimposing the RBDs from crystal structures of SARS-CoV-2 RBD complexed with COVA1-16 (PDB 7JMW) and C022 (this study), the V_H_-V_L_ domains of the bound Fabs were related by an root mean square deviation (RMSD) of 1.3Å (235 Cα atoms), with the majority of conformational differences occurring in the CDRH1 and CDRH2 loops (Figure S5A). Despite being derived from different V gene segments (which would affect their VH gene segment-encoded CDRH1 and CDRH2 loops), C022 and COVA1-16 recognized similar epitopes, contacting a common set of 23 RBD residues, which included COVA1-16 interactions in common with C022 interactions with the RBD α4 helix (Figure 3B).

While C022 and COVA1-16 share a generally similar mode of binding, there are differences in interactions of residues encoded within their different V_H_ gene segments (i.e., their CDRH1 and CDRH2 loops) (Figure S5B). For example, the C022 contact with T430_RBD_ was part of an extensive clasp made by an interaction between the C022 CDRH1 residue R33_HC_ with backbone carbonyls of D427_RBD_, D428_RBD_, and F429_RBD_ and with the sidechain of T430_RBD_ (Figure S5C). Two of the same RBD residues (D427_RBD_ and F429_RBD_) interacted with an arginine from COVA1-16, but this arginine (R100B_HC_) is located at the base of the CDRH3 loop rather than within CDRH1, as is the case with C022 R33_HC_. The larger separation distance from the RBD of COVA1-16 R100B_HC_ allowed it to form a sidechain-backbone H-bond with D427_RBD_ similar to a sidechain–backbone H-bond involving C022 R33_HC_ and D428_RBD_, but precluded interactions with D428_RBD_ and T430_RBD_ (Figure S5D). In addition, the COVA1-16 CDRH1 was shorter than the C022 CDRH1 (7 versus 9 residues) (Figure S4A), and was shifted away from the RBD relative to the C022 CDRH1. These differences, in addition to fewer LC interactions by COVA1-16, resulted in less total BSA for COVA1-16 relative to C022 (1607 Å^2^ vs 1875 Å^2^, respectively) despite similar contributions from CDRH3 loops (Table S2).

## Interactions with RBD main chain atoms facilitate recognition of diverse RBDs

The paratopes of both C118 and C022 were dominated by their long CDRH3 loops (20 and 21 residues, respectively) (Figure 4A,B and Figure S4), which make up ∼half of the buried surface areas (BSAs) of each paratope (461 Å^2^ of 1020 Å^2^ for C118 and 537 Å^2^ of 969 Å^2^ for C022) (Table S2). The C118 and C022 CDRH3s comprise two anti-parallel β-strands that extend a largely internal RBD β-sheet (β-strands β1-β4 and β7) through main chain H-bonds between the RBD β2 strand (377-379_RBD_) and the first CDRH3 β-strand (CDRH3 residues 97-99 (C118) or 100-100B (C022)) (Figure 4A,B). A similar feature is also seen in the structure of the COVA1-16–RBD complex (Liu et al., 2020a), which shares a nearly identical CDRH3 sequence with C022 (Figure S5E).

**Figure 4:**
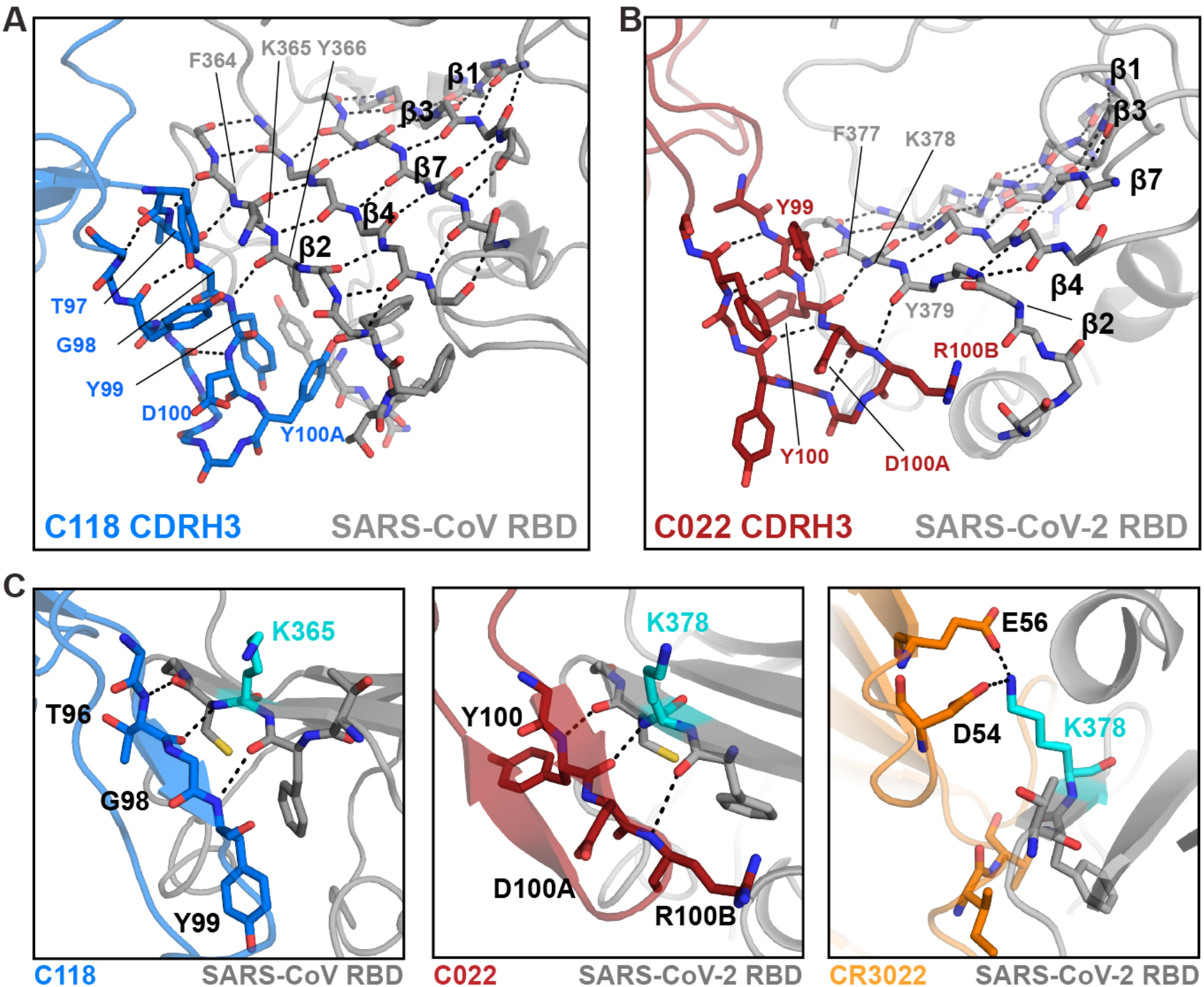
C118 and C022 Fabs primarily use their CDRH3s for contacts with RBD alternative binding contacts for K378_RBD_. (A) Close-up cartoon showing β-hairpin formed by C118 CDRH3 (blue sticks) and β-sheet formation with SARS-CoV RBD (grey cartoon with sticks). H-bonds shown as black dashed lines. (B) Close-up cartoon showing β-hairpin formed by C022 CDRH3 (red sticks) and β-sheet formation with SARS-CoV-2 RBD (grey cartoon with sticks). H-bonds shown as black dashed lines. (C) Cartoon and stick representation of C118-RBD (left), C022-RBD (middle) and CR3022-RBD (right) showing distinct interactions with residue K365_SARS_/K378_SARS2_ (cyan).

C118 and C022 form extensive backbone interactions with RBD, with 10 and 9 H-bonds formed with the backbone of RBD, respectively. Extensive backbone interactions in the C118 and C022 epitopes could contribute to their breadth of binding and neutralization across sarbecoviruses, as backbone interactions would facilitate binding despite side chain substitutions, which are rare across the RBD sequences listed (Figure 3B), but could occur in other CoV RBDs. For example, the backbone H-bonds between the CDRH3s of C118 and C022 with the RBD β2 strand allow for binding despite substitution at position K378_RBD_ (K365_RBD_ in SARS-CoV) (Figure S2A and Figure 4C). By contrast, the class 4 antibody CR3022 uses side chain interactions (potential electrostatic interactions between D54_HC_ and E56_HC_ and K378_RBD_); thus CR3022 is sensitive to mutation at K378N_RBD_ (Figure S2A). This is consistent with CR3022 not binding to Rf1-CoV RBD (Figure 1C), which contains an asparagine at the equivalent position to SARS-CoV-2 K378_RBD_ (Figure 3B), whereas C118 and C022 binding to Rf1-CoV RBD was not affected. Overall, mainchain H-bond interactions likely reduce sensitivity to RBD sidechain substitutions, making antibodies such as C118 and C022 more tolerant to differences between sarbecoviruses strains or variants.

### C118-S cryo-EM structure shows increased S trimer opening

On an S trimer, the class 4 cryptic epitope is at the base of the RBD, where it faces towards the center of the trimer (Barnes et al., 2020a; Huo et al., 2020; Yuan et al., 2020b). The epitope is buried in the closed, prefusion S conformation and interacts with portions of the spike S2 subunit and neighboring ‘down’ RBDs. Compared to class 2 or class 3 anti-RBD antibodies that recognize their epitopes in ‘up’ or ‘down’ RBD conformations (Barnes et al., 2020a), the class 4 epitope is less accessible and requires two ‘up’ RBDs for antibody binding (Piccoli et al., 2020b). Additionally, class 4 antibody binding may also require RBD rotation to prevent steric clashes with neighboring ‘up’ RBDs, as observed for the complexes of S trimer with EY6A, S2A4, and S304 (Piccoli et al., 2020b; Zhou et al., 2020a).

Given the similar binding poses of C118 and C022 antibodies, which bind with a more acute angle with respect to the RBD than EY6A or CR3022 (Figure 2E), and the increased breadth and potency of C118 and C022 relative to other class 4 anti-RBD antibodies (Figure 1D), we sought to understand the requirements for epitope recognition on a S trimer. Thus, we solved a single-particle cryo-EM structure of C118 Fabs bound to SARS-CoV-2 S 6P trimers (Hsieh et al., 2020), finding two distinct states defined by RBDs adopting various rotational conformations (Figure 5A,B and Figure S6), as well as C118 Fab bound to dissociated S1 subunit protomers (Figure S6B). For the state 1 C118-S trimer complex structure solved to 3.4Å, we subsequently used symmetry expansion and local refinement to generate a 3.7Å map of the C118 V_H_V_L_ – RBD interface (Figure S6B-E).

**Figure 5.**
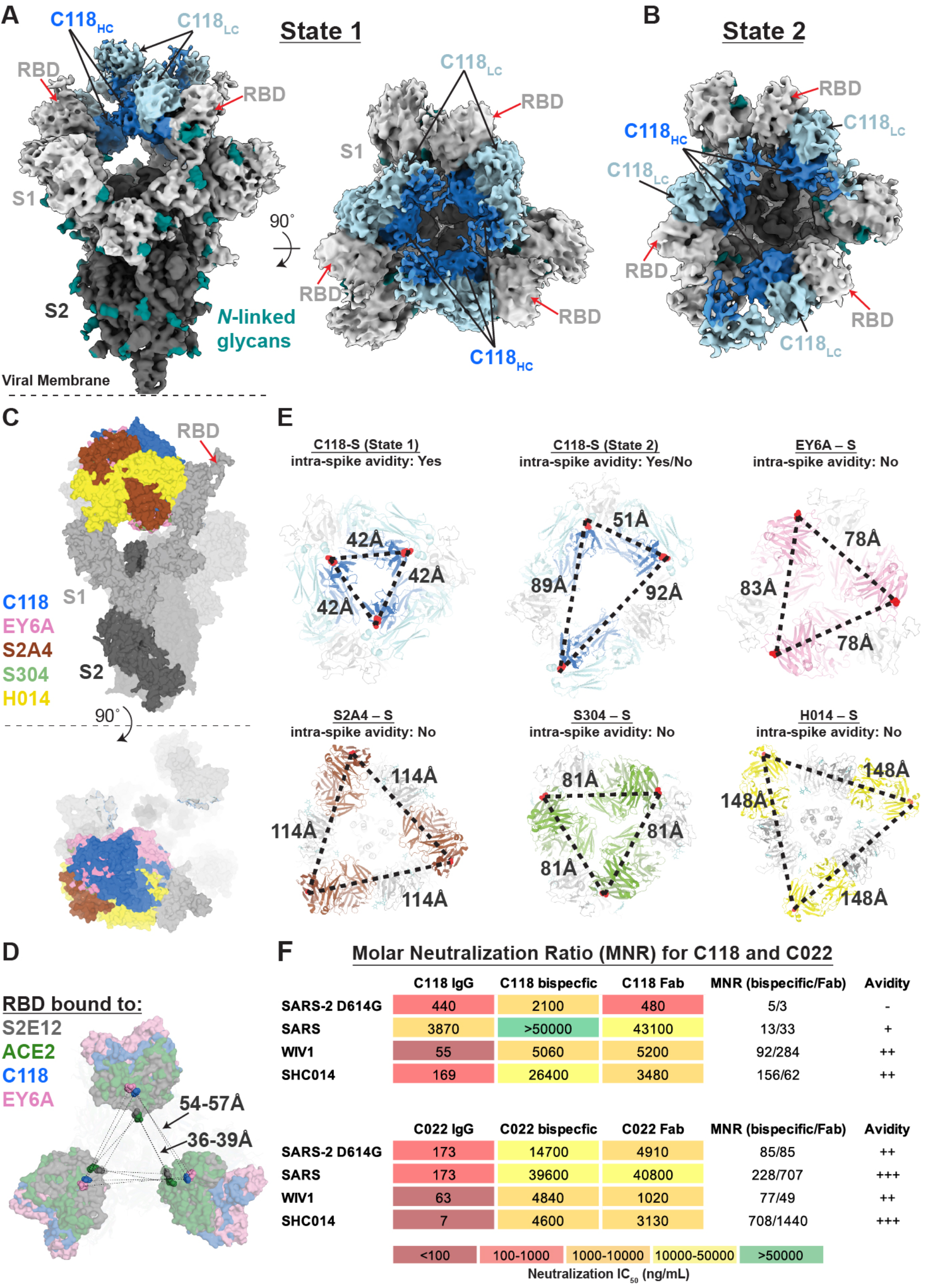
Cryo-EM structure of C118-S complex shows binding to cryptic epitope and the potential for intra-spike crosslinking. (A) 3.4Å cryo-EM density for the C118 – S trimer complex (State 1). Side view (left panel) illustrates orientation with respect to the viral membrane (dashed line). Top view (right panel) shows symmetric binding at the trimer apex with C118 HC (blue) oriented in the interior. (B) 4.4Å cryo-EM density for the C118 – S trimer complex (State 2). Top view illustrates asymmetry of complex due to RBD rotation in one protomer. (C) Composite model of an open SARS-CoV-2 trimer bound by class 4 Fabs: C118 (this paper, blue), EY6A (PDB 6ZDH, pink), S2A4 (PDB 7JVC, brown), the class 4 anti-SARS antibody S304 (PDB 7JW0, green), and H014 (PDB 7CAK, yellow). (D) Comparison of S trimer openness by measurements of Cα distances for D428_RBD_ between adjacent ‘up’ RBDs in S trimers complexed with: the class 1 antibody S2E12 (PDB 7K43, gray), soluble ACE2 (PDB 7KMS, green) and the class 4 antibodies C118 (this study, blue) and EY6A (PDB 6ZDH, pink). (E) Prediction of potential intra-spike avidity effects by measurement of Cα distances between the C-termini of adjacent C_H_1 domains for the mAb-S trimer complexes described in panel C. Measurements were used to evaluate the potential for intra-spike crosslinking by an IgG binding to a single spike trimer as described (Barnes et al., 2020a). For the H014-S complex, the C_H_1-C_L_ domains were rigid body fit into the cryo-EM density (EMD-30333) prior to measurements. (F) IC_50_ values and molar neutralization ratios (MNRs) defined as: [IC_50_ Fab or bispecific IgG (nM)/IC_50_ IgG (nM)] (Klein and Bjorkman, 2010) for C118 and C022. IC_50_ values shown for the IgGs are from Figure 1D. IC_50_ values for all assays against SARS-CoV-2 and SARS-CoV are means of 2-7 independent experiments. Two MNRs are presented in the MNRs (bispecific/Fab) column: the MNR calculated using a bispecific IgG versus the bivalent IgG (left) and the MNR calculated using a Fab versus the bivalent IgG (right). Neutralization results with MNRs ≤5 are indicated as not demonstrating avidity effects (-), >10 are indicated as demonstrating minimal avidity (+), results with one MNR> 50 are indicated as moderate avidity (++), and MNRs demonstrating strong avidity effects (one MNR >700) are indicated as +++.

The C118 pose with respect to the RBD observed in the C118 – SARS-CoV-2 S structure was similar to the C118 – SARS-CoV RBD crystal structure (Figure S6F), demonstrating consistent recognition of the antibody epitope on both SARS-CoV-2 and SARS-CoV RBDs. Furthermore, the C118 binding pose was oriented higher on the RBD relative to other class 4 anti-RBD antibodies (Figure 5C), and was consistent with SPR competition data that suggested C118 would sterically hinder ACE2 binding to the same protomer (Figure S3).

Despite differences in binding poses relative to other class 4 antibodies (Figure 5C), C118 binding also resulted in RBD conformations displaced further from the trimer center relative to S2E12 (a class 1 anti-RBD neutralizing antibody) (Tortorici, 2020) and ACE2 (Yan et al., 2020) (Figure 5D). On average, class 4 anti-RBD antibody binding resulted in an ∼15-20Å displacement of the RBD relative to ACE2-bound conformations, which likely results in destabilization of the spike trimer. Indeed, S1 shedding induced by class 4 antibodies has been described as a possible neutralization mechanism (Huo et al., 2020; Piccoli et al., 2020b; Wec et al., 2020). The presence of C118-S1 protomer classes in our cryo-EM data suggested that C118 also induces shedding (Figure S6), but the role S1 shedding and premature S-triggering plays in C118-mediated neutralization requires further investigation.

### C118 and C022 neutralization of sarbecoviruses demonstrate differential effects of avidity enhancement

Neutralization of SARS-CoV-2 and SARS-CoV by COVA1-16 was found to be mediated by avidity effects based on potent neutralization by the bivalent COVA1-16 IgG, but not the monovalent Fab (Liu et al., 2020a). To evaluate whether intra-spike crosslinking, one source of avidity enhancement for bivalent antibodies, was possible for C118 or C022 IgGs, we examined the C118-S trimer structure to ask whether the positioning of two Fabs on adjacent RBDs would be compatible with binding by a single IgG. As previously described for other anti-RBD IgGs, we compared the distance between residues near the C-termini of adjacent Fab C_H_1 domains to analogous distances in crystal structures of intact IgGs, setting a cut-off of ≤65Å as potentially allowing a single IgG to include both Fabs (Barnes et al., 2020a). The measured distance for the C-termini of adjacent Fab C_H_1 domains in the symmetric State 1 C118-S trimer structure was 41Å (Figure 5E), suggesting that intra-spike crosslinking would be possible for C118 IgGs bound to spike trimers. The asymmetric State 2 C118-S structure included distances of 50Å, 89Å, and 92Å (Figure 5E), also allowing intra-spike crosslinking between one combination of two bound RBDs, as well as the potential for inter-spike crosslinking between adjacent spikes on the virion surface. In comparison, no other class 4 anti-RBD Fab-S trimer structures showed measured distances that would be compatible with intra-spike crosslinking (Figure 5E), thus any potential avidity effects for those IgGs could only occur via inter-spike crosslinking.

To further evaluate whether avidity could also facilitate cross-reactive neutralization by the C118 and C022 antibodies, we compared neutralization of SARS-CoV-2, SARS-CoV, WIV1, and SHC014 by the bivalent C118 and C022 IgGs and by two monovalent forms of each antibody: a 50 kDa Fab and an IgG size-matched bispecific IgG containing only one relevant Fab. The bispecific IgGs included one C118 or C022 RBD-binding Fab and a second non-RBD-binding Fab derived from the HIV-1 antibody 3BNC117 (Scheid et al., 2011). To interpret neutralization results, we calculated molar neutralization ratios (MNRs) defined as: [IC_50_ Fab or bispecific IgG (nM)/IC_50_ IgG (nM)] (Klein and Bjorkman, 2010). In the absence of avidity effects resulting from either crosslinking within a spike trimer (intra-spike crosslinking) or cross-linking between adjacent spike trimers (inter-spike crosslinking), an MNR would be 2.0, which accounts for twice as many relevant Fabs in a bivalent IgG compared to its monovalent forms.

Using pseudotyped SARS-CoV-2, SARS-CoV, WIV1, and SHC014, we derived neutralization potencies of the bivalent IgG, monovalent bispecific IgG, and Fab forms of C118 and C022 and then calculated MNRs for the bivalent IgG to bispecific IgG comparison (bispecific MNR) and for the bivalent IgG to Fab comparison (Fab MNR) (Figure 5F). Comparisons between the Fab and bispecific IgG forms of monovalent antibody allowed evaluation of potential steric affects that could increase neutralization potencies for larger IgGs compared to smaller Fabs. With the exception of the low MNRs derived from the IgG comparison with the bispecific and Fab forms of C118 against SARS-CoV-2 (MNRs of 5 and 3, respectively), we found mostly high MNRs ranging from the lowest values of >11 and 28 for the MNRs for C118 against SARS-CoV (where 11 is a minimal estimate since the C118 bispecific was non-neutralizing) to the highest values of 708 and 1444 for the C022 bispecific and Fab MNRs against SHC014. Four of the bispecific to Fab MNR comparisons showed a two-fold or higher Fab MNR than the comparable bispecific MNR, suggesting that at least some of the increased potencies of the bivalent IgGs compared with their counterpart Fabs resulted from steric effects. However, six of the eight monovalent to bivalent comparisons exhibited MNRs well over 70, suggestive of strong avidity effects. By contrast, mean MNRs derived for broadly neutralizing anti-HIV-1 Env antibodies are ≤10 (Wang et al., 2017), consistent with the low spike density on HIV-1 virions that largely prevents inter-spike crosslinking, and the architecture of the HIV-1 Env trimer, which prohibits intra-spike crosslinking for all known HIV-1 broadly neutralizing antibodies (Klein and Bjorkman, 2010). Taken together with the analysis of the C118-S trimer structure, the observed avidity effects for C118 IgGs binding to WIV1 and SHC014 and for the related C022 IgGs binding to the four viruses tested could arise from intra-spike as well as inter-spike crosslinking.

The question as to why C118 exhibits little or no avidity effects for neutralization of SARS-CoV-2 and SARS-CoV is difficult to address since the same IgG showed strong avidity effects against WIV1 and SHC014, and C022, which binds similarly to C118, showed avidity effects in neutralization of all four pseudoviruses. These results could derive from different binding characteristics for C118 to the SARS-CoV-2 and SARS-CoV RBDs compared with C118 and C022 interactions with the other sarbecoviruses evaluated. Indeed, simulations of avidity effects demonstrated that some combinations of IgG concentration and antigen-binding affinity and kinetic constants showed no advantages of bivalent versus monovalent binding (Klein, 2009; Klein and Bjorkman, 2010). Thus the effects of avidity are a complicated function of concentration and binding constants that preclude predictions in the absence of experimental data.

## Discussion

Concerns about coronaviruses having spillover potential as well as the increasing prevalence of SARS-CoV-2 variants necessitates identification of cross-reactive antibodies. Antibodies elicited against infectious viruses for which there are multiple circulating variants, either within an individual or the population, often show a trade-off between potency and breadth (Corti et al., 2010; Desrosiers et al., 2016). In the case of antibody responses against SARS-CoV-2, the cause of the current global pandemic, many strongly neutralizing antibodies have been isolated that block ACE2 receptor interactions (Barnes et al., 2020a; Dejnirattisai et al., 2021; Lee et al., 2021; Liu et al., 2020b; Piccoli et al., 2020a). However, the ACE2-binding region of the RBD also tends to accumulate amino acid changes, as evidenced by substitutions identified in the current SARS-CoV-2 variants of concern (Annavajhala et al., 2021; Faria et al., 2021; Rambaut et al., 2020; Tegally et al., 2020; Voloch et al., 2020; West et al., 2021; Zhang et al., 2021), thus reducing the potential efficacies of vaccines and monoclonal antibody therapies. Recent studies suggest that antibodies against the S2 subunit offer the potential of greater cross-reactivity across coronaviruses, but these antibodies generally lack strong neutralization potency (Sauer et al., 2021; Shah et al., 2021; Wang et al., 2021).

The class 4 RBD-binding epitope, which is more conserved than the class 1 and class 2 RBD epitopes, represents a plausible target for the elicitation of antibodies with broad cross-reactive recognition across sarbecoviruses. Indeed, some recently described class 4 antibodies (e.g., CR3022, H014, COVA1-16, EY6A, ADI-56046) neutralize two or more sarbecovirus strains, and/or can bind RBDs from multiple sarbecoviruses (Liu et al., 2020a). However, while many class 4 antibodies show some cross-reactivity, they generally exhibit decreased potencies against heterologous sarbecovirus strains. For example, the SARS-CoV–derived CR3022 antibody does not potently neutralize SARS-COV-2 (Huo et al., 2020), and the SARS-CoV-2 – derived COVA1-16 antibody does not potently neutralize SARS-CoV or SARS-CoV-2 variants of concern (Liu et al., 2020a; Liu et al., 2021).

Here we characterized two antibodies, C118 and C022, derived from different COVID-19 convalescent donors (Robbiani et al., 2020), which show breadth of and potent neutralization against sarbecoviruses of all three clades. The structural similarity of RBD binding poses between C022 and COVA1-16 (Liu et al., 2020a), which was derived from yet a third COVID-19 convalescent donor (Brouwer et al., 2020), suggests that these sorts of cross-reactive antibodies are commonly elicited by natural infection and that their epitope represents an attractive target for immunogen design. Of particular importance for the current pandemic, circulating variants of concern or variants of interest did not confer resistance to the C118 and C022 antibodies. In addition, C118 and C022 antibodies were not affected by naturally-occurring RBD mutations that undermine the activity of several antibodies approved for therapeutic use (Hoffmann et al., 2021; Starr et al., 2021b).

Analysis of our C118-RBD and C022-RBD complex structures revealed key details of cross-reactive recognition and broad sarbecovirus neutralization. First, C118 and C022 utilize long CDRH3s to facilitate interactions with the cryptic RBD epitope at the base of the RBD. In contrast to less potent class 4 antibodies such as CR3022 (Huo et al., 2020; Yuan et al., 2020a; Yuan et al., 2020b) and EY6A (Zhou et al., 2020a) that also contact this region, the longer CDRH3 provides the opportunity to target a highly-conserved patch of residues across sarbecoviruses with an orientation that extends the epitope upwards to the ACE2 binding site, a structural feature shared with COVA1-16 (Liu et al., 2020a). Second, the aforementioned binding poses of C118, C022, and COVA1-16, as well as overlap of the C022 epitope with the edge of the ACE2 binding site, suggested competition with ACE2 as part of their neutralization mechanisms. Indeed, competition experiments reported here for C118 and C022 and by others for COVA1-16 (Liu et al., 2020a) demonstrated competition with ACE2 for SARS-CoV-2 RBD binding. Third, C118 and C022 formed many interactions with backbone atoms of RBD residues, adding a second level of buffering against viral escape since amino acid substitutions at these positions are less likely to abrogate antibody binding. Finally, the demonstration that C118 and C022 bivalency increased potency of neutralization against some of the viruses evaluated showed the potential for these antibodies to utilize avidity effects for neutralization of sarbecoviruses. Given the requirement for two ‘up’ RBDs on a S trimer for class 4 antibody binding, bivalent binding within a single S trimer would be possible. Thus we suggest that intra-spike crosslinking would be an advantage for neutralization of sarbecoviruses, where avidity effects likely play a role.

In conclusion, class 4 antibodies that access the cryptic RBD epitope and compete with ACE2 binding are important for understanding cross-reactivity of human SARS-CoV-2 antibody responses elicited by natural sarbecovirus infection. We suggest that potent class 4 anti-RBD antibodies could be used therapeutically to avoid resistance to SARS-CoV-2 variants of concern, perhaps after in vitro selection to further improve their potencies. Structural characterization of these antibodies could also be used to inform future immunogen design efforts to elicit cross-reactive antibodies against SARS-CoV-2 variants and other sarbecoviruses.

## Acknowledgements

We thank J. Vielmetter, P. Hoffman, and the Protein Expression Center in the Beckman Institute at Caltech for expression assistance and K. Dam for assistance with soluble ACE2 purification. Electron microscopy was performed in the Caltech Cryo-EM Center with assistance from S. Chen and A. Malyutin. We thank the Gordon and Betty Moore and Beckman Foundations for gifts to Caltech to support the Molecular Observatory. We thank J. Kaiser, director of the Molecular Observatory at Caltech, and beamline staff C. Smith and S. Russi at SSRL for data collection assistance. Use of the Stanford Synchrotron Radiation Lightsource, SLAC National Accelerator Laboratory, is supported by the U.S. Department of Energy, Office of Science, Office of Basic Energy Sciences under Contract No. DE-AC02-76SF00515. The SSRL Structural Molecular Biology Program is supported by the DOE Office of Biological and Environmental Research, and by the National Institutes of Health, National Institute of General Medical Sciences (P30GM133894). The contents of this publication are solely the responsibility of the authors and do not necessarily represent the official views of NIGMS or NIH. This work was supported by NIH (P01-AI138938-S1 to P.J.B. and M.C.N.), the Caltech Merkin Institute for Translational Research (P.J.B.), a George Mason University Fast Grant (P.J.B.), NIH R01AI078788 (to T.H.) and R01AI640511 (to P.D.B). C.O.B was supported by the Hanna Gray Fellowship Program from the Howard Hughes Medical Institute and the Postdoctoral Enrichment Program from the Burroughs Wellcome Fund. M.C.N. is an HHMI investigator.

## Author Contributions

C.A.J., A.A.C., P.J.B., and C.O.B. conceived and designed experiments. Proteins were produced and characterized by A.A.C., K.H.T., C.O.B., and C.A.J. Binding and neutralization studies were done by A.A.C., and F.M. with assistance from P.N.P.G., Y.L., and F.S. SPR binding competition experiments were done by C.A.J. with assistance from J.R.K. Structural studies were performed by C.A.J. with assistance from C.O.B. Structure analysis was done by C.A.J. with assistance from C.O.B. and A.A.C. Sequence analysis was done by A.P.W. Paper was written by C.A.J., P.J.B., and C.O.B. with assistance from A.A.C., T.H., M.C.N., P.D.B., and other authors.

## STAR Methods

### RESOURCE AVAILABILITY

#### Lead Contact

All requests for further information or reagents should be directed to the Lead Contact, Pamela Bjorkman.

#### Materials Availability

All expression plasmids generated in this study for human CoV proteins, Fabs and IgGs are available upon request.

#### Data and Code Availability

Atomic models of C118 Fab complexed with SARS-CoV RBD and C022 Fab complexed with SARS-CoV-2 RBD have been deposited in the Protein Data Bank (PDB) (http://www. rcsb.org/) under accession codes XXXX and YYYY, respectively. The atomic model and cryo-EM map generated for the C118 Fab–SARS-CoV-2 S complex have been deposited at the PDB (http://www.rcsb.org/) and the Electron Microscopy Databank (EMDB) (http://www.emdataresource.org/) under accession codes AAAA and EMD-BBBBB, respectively.

### EXPERIMENTAL MODEL DETAILS

#### Cell lines

Cells for pseudovirus production (HEK293T) were cultured at 37°C and 5% CO_2_ in Dulbecco’s modified Eagle’s medium (DMEM, Gibco) supplemented with 10% heat-inactivated fetal bovine serum (FBS, Sigma-Aldrich) and 5 µg/ml Gentamicin (Sigma-Aldrich).

Target cells for pseudovirus neutralization experiments (HEK293T_ACE2_) were generated as described (Robbiani et al., 2020) and cultured at 37°C and 5% CO_2_ in Dulbecco’s modified Eagle’s medium (DMEM, Gibco) supplemented with 10% heat-inactivated fetal bovine serum (FBS, Sigma-Aldrich), 5 µg/ml gentamicin (Sigma-Aldrich), and 5µg/mL Blasticidin (Gibco).

Expi293F cells (Gibco) for protein expression were maintained at 37°C and 8% CO_2_ in Expi293 expression medium (Gibco), transfected using an Expi293 Expression System Kit (Gibco) and maintained under shaking at 130 rpm. All cell lines were female and were not specifically authenticated.

#### Bacteria

*E. coli* DH5 Alpha (Zymo Research) used for propagation of expression plasmids were cultured with shaking at 250 rpm at 37°C in LB broth (Sigma-Aldrich).

#### Viruses

To generate pseudotyped viral stocks, HEK293T cells were transfected with pNL4-3ΔEnv-nanoluc and pSARS-CoV2-S_trunc_ (Robbiani et al., 2020) using polyethylenimine, leading to production of HIV-1-based pseudovirions carrying the SARS-CoV-2 S protein at the surface. Eight hours after transfection, cells were washed twice with phosphate buffered saline (PBS) and fresh media was added. Supernatants containing pseudovirus were harvested 48 hours post transfection, filtered and stored at −80°C. Infectivity of pseudoviruses was determined by titration on 293T_ACE2_ cells.

### METHOD DETAILS

#### Phylogenetic trees

Sequence alignments of RBDs were made with Clustal Omega (Sievers et al., 2011). Phylogenetic trees were calculated from amino acid alignments using PhyML 3.0 (Guindon et al., 2010) and visualized with PRESTO (http://www.atgc-montpellier.fr/presto).

#### Protein Expression

Fabs and IgGs were expressed and purified as previously described (Scharf et al., 2015; Schoofs et al., 2019) and stored at 4 °C. Bispecific IgGs (C118 or C022 plus 3BNC117, a non-coronavirus binding HIV-1 antibody (Scheid et al., 2011)) were produced by co-transfection of two heavy chain and two light chain genes that included knobs-into-holes mutations in IgG Fc and a domain cross-over in the 3BNC117 Fab to prevent incorrect light chain pairing (Schaefer et al., 2011). Antibody CDR lengths were determined using the IMGT definitions (Lefranc et al., 2015; Lefranc et al., 2009).

The following C-terminally 6xHis-tagged RBD proteins were transfected and expressed as described previously (Cohen et al., 2021): SARS-CoV-2 RBD (residues 328-533), SARS-CoV-2 RBD mutants (residues 319-541), SARS RBD (residues 318-510), SHC014 RBD (residues 307-524), WIV-1 RBD (residues 307-528), RaTG13 RBD (residues 319-541), Rs4081 RBD (residues 310-515), Yun11 RBD (residues 310-515), Rf1 RBD (residues 310-515), RmYN02 RBD (298-503), BM-4831 RBD (residues 310-530), BtKY72 RBD (residues 309-530). A trimeric SARS-CoV-2 ectodomain (residues 16-1206 of the early SARS-CoV-2 GenBank MN985325.1 sequence isolate with 6P (Hsieh et al., 2020) stabilizing mutations, a mutated furin cleavage site between S1 and S2, a C-terminal TEV site, foldon trimerization motif, octa-His tag, and AviTag) was expressed as described (Barnes et al., 2020a; Barnes et al., 2020b). A gene encoding a 6xHis-tagged soluble human ACE2 construct (residues 1-615) was purchased from Addgene (Catalog # 149268) and expressed and purified as described (Chan et al., 2020).

SARS-CoV-2 S trimer, RBDs, and soluble ACE2 were purified by Nickel-NTA and size-exclusion chromatography using a Superdex 200 column (GE Life Sciences) as described (Barnes et al., 2020a; Cohen et al., 2021). Peak fractions were identified by SDS-PAGE, and those containing S trimer, monomeric RBDs, or soluble ACE2 were pooled, concentrated, and stored at 4 °C (RBDs) or flash frozen in nitrogen and stored at −80 °C (S trimer) until use.

#### ELISAs

Purified RBD at 10 µg/ml in 0.1 M NaHCO_3_ pH 9.8 was coated onto Nunc® MaxiSorp™ 384-well plates (Sigma) and stored overnight at 4°C. The following day, plates were blocked with 3% bovine serum albumin (BSA) in TBS-T Buffer (TBS + 0.1% Tween20) for 1hr at room temperature. Blocking solution was removed from the plates, purified IgGs at 50 µg/mL were serially diluted by 4-fold with TBS-T/3% BSA and added to plates for 3 hr at room temperature. Plates were washed with TBS-T and then incubated with 1:15,000 dilution of secondary HRP-conjugated goat anti-human IgG for 45 minutes at room temperature (Southern Biotech). Plates were washed again with TBS-T and developed using SuperSignal™ ELISA Femto Maximum Sensitivity Substrate (ThermoFisher) and read at 425 nm. ELISAs were done in duplicate, and curves were plotted and integrated to obtain the area under the curve (AUC) using Graphpad Prism v9.1.0.

#### Neutralization assays

SARS-CoV-2, SARS-CoV-2 variants of concern (Annavajhala et al., 2021; Faria et al., 2021; Rambaut et al., 2020; Tegally et al., 2020; Voloch et al., 2020; West et al., 2021; Zhang et al., 2021), SARS-CoV, WIV1, and SHC014 pseudoviruses based on HIV-1 lentiviral particles were prepared as described (Cohen et al., 2021; Crawford et al., 2020; Robbiani et al., 2020) using genes encoding S protein sequences with cytoplasmic tail deletions: 21 amino acid deletions for SARS-CoV-2, SARS-CoV-2 variants of concern, WIV1, and SHC014 and a 19 amino acid deletion for SARS-CoV. Plasmids expressing the spike protein found in the bat (*Rinolophus Sinicus*) coronavirus bCoV-WIV16 as well as the pangolin (*Manis javanica*) coronaviruses from Guandong, China (pCoV-GD) and Guanxi, China (pCoV-GX) have been described previously and are based on ALK02457 (Genebank), Pangolin_CoV_EPI_ISL_410721(Gisaid) and Pangolin_CoV_EPI_ISL_410542 (Gisaid) (Muecksch et al., 2021).

Relative to the SARS-CoV-2 spike gene (Wuhan-Hu-1 Spike Glycoprotein Gene, D614G mutant, designated as ‘wt’ in Figure 1D), the SARS-CoV-2 variants of concern included the D614G mutation and the following other substitutions: B.1.351: L18F, D80A, D215G, del242-244, R246I, K417N, E484K, N501Y, A701V; B.1.1.7: del69-70, del144, N501Y, A570D, P681H, T716I, S982A, D1118H; B.1.429: S13I, W152C, L452R, and B.1.526: L5F, T95I, D253G, E484K, A701V. For neutralization assays presented in Figure 1D, four-fold dilutions of purified IgGs (starting concentrations of 50 µg/mL) were incubated with a pseudotyped virus for 1 hour at 37°C. Cells were washed twice with phosphate-buffered saline (PBS) and lysed with Luciferase Cell Culture Lysis 5x reagent (Promega) after incubation with 293T_ACE2_ target cells for 48 hours at 37°C. NanoLuc Luciferase activity in lysates was measured using the Nano-Glo Luciferase Assay System (Promega). Relative luminescence units (RLUs) were normalized to values derived from cells infected with pseudotyped virus in the absence of IgG. Half-maximal inhibitory concentrations (IC_50_ values) were determined using 4- or 5-parameter nonlinear regression in AntibodyDatabase (West et al., 2013).

Relative to the SARS-CoV-2 spike gene (Wuhan-Hu-1; NC_045512, D614 sequence designated as ‘wt’ in Figure S2), a panel of plasmids expressing RBD mutant SARS-CoV-2 S proteins in the context of pSARS-CoV-2-S_Δ19_ have been described previously (Muecksch et al., 2021; Robbiani et al., 2020; Schmidt et al., 2020; Weisblum et al., 2020). The E484K substitution was constructed in the context of a pSARS-CoV-2-S_Δ19_ variant with a mutation in the furin cleavage site (R683G) to increase infectivity (Muecksch et al., 2021). The IC_50_ values of this pseudotype (E484K/R683G) was compared to a wild-type SARS-CoV-2 S sequence carrying R683G in the subsequent analyses. For neutralization assays presented in Figure S2, monoclonal antibodies were four-fold serially diluted and incubated with SARS-CoV-2 pseudotyped HIV-1 reporter virus for 1 h at 37 °C (final starting concentration of 2.5 µg/ml). The antibody and pseudotyped virus mixture was added to HT1080ACE2.cl 14 cells (Schmidt et al., 2020). After 48 h, cells were washed with PBS and lysed with Luciferase Cell Culture Lysis 5× reagent (Promega). Nanoluc luciferase activity in cell lysates was measured using the Nano-Glo Luciferase Assay System (Promega) and the Glomax Navigator (Promega). Relative luminescence units were normalized to those derived from cells infected with SARS-CoV-2 pseudovirus in the absence of monoclonal antibodies. The 50% inhibitory concentration (IC_50_) was determined using 4-parameter nonlinear regression (least-squares regression method without weighting; constraints: top = 1, bottom = 0) (GraphPad Prism).

#### SPR-based ACE2 binding competition experiments

Surface Plasmon Resonance (SPR) experiments were done using a Biacore T200 instrument (GE Healthcare).

Surface Plasmon Resonance (SPR) experiments were done using a Biacore T200 instrument (GE Healthcare). Purified SARS CoV-2 RBD was conjugated to each of the four flow cells using primary amine chemistry at pH 4.5 (Biacore manual) to a CM5 chip (GE Healthcare) to a response level of ∼700 resonance units (RUs). C118, C022, C144, and CR3022 IgG (1000nM) in buffer HBS-EP+ (150mM sodium chloride, 10mM HEPES, 3mM EDTA, 0.05% Tween-20, pH 7.6) were each injected on the RBD-CM5 chip for a contact time of 600 sec at 30µL/min. A second injection of soluble ACE2 at 250nM was injected over the immobilized RBD-Fab at 30µL/min for a contact time of 300 sec and dissociation time of 30 sec in HBS-EP+ buffer. Data were analyzed and plotted using Prism 9 (Graphpad).

#### X-ray crystallography

Fab-RBD complexes were assembled by incubating an RBD with a 1.5x molar excess of Fab for 1 hr on ice followed by size exclusion chromatography on an S200 10/300 increase column (GE Life Sciences). Fractions containing complex were pooled and concentrated to 8mg/mL. Crystallization trials using commercially-available screens (Hampton Research) were performed at room temperature using the sitting drop vapor diffusion method by mixing equal volumes of a Fab-RBD complex and reservoir using a TTP LabTech Mosquito instrument. Crystals were obtained for C118 Fab-SARS RBD in 0.2M sodium fluoride, 20% w/v polyethylene glycol 3,350 and for C022 Fab-SARS-CoV-2 RBD in 0.05M ammonium sulfate, 0.05M Bis-Tris, 30% v/v pentaerythritol ethoxylate (15/4 EO/OH). Crystals were cryoprotected by adding glycerol directly to drops to a final concentration of 20% and then looped and cryopreserved in liquid nitrogen.

X-ray diffraction data were collected at the Stanford Synchrotron Radiation Lightsource (SSRL) beamline 12-2 on a Pilatus 6M pixel detector (Dectris). Data from single crystals were indexed and integrated in XDS (Kabsch, 2010) and merged using AIMLESS in *CCP4* (Winn et al., 2011) (Table S1). The C022-RBD structure was solved by molecular replacement in PHASER (McCoy et al., 2007) using unmodified RBD coordinates (PDB 7K8M) and coordinates from C102 Fab (PDB 7K8M) after trimming heavy chain and light chain variable domains using Sculptor (Bunkóczi and Read, 2011) as search models. Coordinates were refined with *phenix.refine* from the PHENIX package ver. 1.17.1 (Adams et al., 2010) and cycles of manual building in Coot (ver 0.8.9.1) (Emsley et al., 2010) (Table S1).

#### Cryo-EM Sample Preparation

C118 Fab-S trimer complex was assembled by incubating purified SARS-CoV-2 S trimer at a 1.2:1 molar excess of purified Fab per S protomer at RT for 30 min. 17 uL of complex was mixed with 0.8uL of a 0.5% w/v F-octylmaltoside solution (Anatrace) and then 3µL were immediately applied to a 300 mesh, 1.2/1.3 AuUltraFoil grid (Electron Microscopy Sciences) that had been freshly glow discharged for 1 min at 20mA using a PELCO easiGLOW (Ted Pella). The grid was blotted for 3.5s with Whatman No. 1 filter paper at 22°C and 100% humidity then vitrified in 100% liquid ethane using a Mark IV Vitrobot (FEI) and stored under liquid nitrogen.

#### Cryo-EM data collection and processing

Single-particle cryo-EM data were collected for the C118-S trimer complex as previously described (Barnes et al., 2020a). Briefly, for the C118-S trimer complex, micrographs were collected on a Talos Arctica transmission electron microscope (Thermo Fisher) operating at 200 kV using a 3×3 beam image shift pattern with SerialEM automated data collection software (Mastronarde, 2005). Movies were obtained on a Gatan K3 Summit direct electron detector operating in counting mode at a nominal magnification of 45,000x (super-resolution 0.4345 Å/pixel) using a defocus range of −0.7 to −2.0 μm. Movies were collected with an 3.6 s exposure time with a rate of 13.5 e^-^/pix/s, which resulted in a total dose of ∼60 e-/Å^2^ over 40 frames. The 2,970 movies were patch motion corrected for beam-induced motion including dose-weighting within cryoSPARC v3.1 (Punjani et al., 2017) after binning super resolution movies by 2 (0.869 Å/pixel). The non-dose-weighted images were used to estimate CTF parameters using Patch CTF in cryoSPARC, and micrographs with poor CTF fits and signs of crystalline ice were discarded, leaving 2,487 micrographs. Particles were picked in a reference-free manner using Gaussian blob picker in cryoSPARC (Punjani et al., 2017). An initial 923,707 particle stack was extracted, binned x4 (3.48 Å/pixel), and subjected to *ab initio* volume generation (4 classes) and subsequent heterogeneous refinement. The 3D classes that showed features for a Fab-S trimer complex were 2D classified to identify class averages corresponding to intact S-trimer complexes with well-defined structural features. This routine resulted in a new particle stack of 110,789 particles, which were unbinned (0.836 Å/pixel) and re-extracted using a 432 box size. Particles were then moved to Relion v3.1 (Zivanov et al., 2018), for further 3D classification (k=6)., which revealed two distinct states of the C118-S trimer complex.

Particles from state 1 (53,728 particles) and state 2 (31,422 particles) were separately refined using non-uniform 3D refinement imposing either C3 or C1 symmetry in cryoSPARC, respectively, to final resolutions of 3.4 Å and 4.5 Å according to the gold-standard FSC (Bell et al., 2016), respectively. To improve features at the C118-RBD interface, particles from State 1 were symmetry expanded and classified for a focused, non-uniform 3D local refinement in cryoSPARC. A soft mask was generated around the C118 V_H_V_L_ – RBD domains (5-pixel extension, 10-pixel soft cosine edge) for local refinements. These efforts resulted in a modest improvement in the RBD-C118 Fab interface (Figure S6B), with an overall resolution of 3.7 Å according to the gold-standard FSC.

#### Cryo-EM Structure Modeling and Refinement

Initial coordinates were generated by rigid-body docking reference structures into cryo-EM density using UCSF Chimera (Goddard et al., 2007). The following coordinates were used: SARS-CoV-2 S 6P trimer: PDB 7K4N (mutated to include 6P mutations), PDB 7BZ5, and C118 Fab variable domains: this study. These initial models were then refined into cryo-EM maps using one round of rigid body refinement, morphing and real space refinement in Phenix (Adams et al., 2010). Sequence-updated models were built manually in Coot (Emsley et al., 2010) and then refined using iterative rounds of refinement in Coot and Phenix (Adams et al., 2010). Glycans were modeled at possible N-linked glycosylation sites (PNGSs) in Coot using ‘blurred’ maps processed with a variety of B-factors (Terwilliger et al., 2018). Validation of model coordinates was performed using MolProbity (Chen et al., 2010) and is reported in Table S3.

#### Structure Analyses

Interacting residues were determined using PDBePISA (Krissinel and Henrick, 2007) for the C118 and C022 epitopes using the following criteria: Potential H-bonds were assigned using a distance of <3.6A ° and an A-D-H angle of >90°, and the maximum distance allowed for a van der Waals interaction was 4.0A °. H-bonds assigned for the C022-RBD complex should be considered tentative due to the relatively low resolution of the structure (3.2Å). Epitope patches for other antibodies in Figure 4A were defined as residues containing an atom within 4Å of the partner protein as determined in PyMOL (Schrödinger, 2011). Buried surface areas (BSAs) were determined with PDBePISA (Krissinel and Henrick, 2007) using a 1.4A ° probe. Structure figures were made using PyMOL ver. 2.3.5 (Schrödinger, 2011) or UCSF Chimera ver. 1.14 (Goddard et al., 2018). Fab-RBD-ACE2 complex figures (Figure 2E) were made by aligning RBD Cα atoms of Fab-RBD (this study and PDBs 6W41 and 6ZCZ) and RBD-ACE2 structures (PDB 6M0J). As density at position N357_RBD_ for our C118-SARS RBD structure precluded building of the glycan, it was modeled (Figure 2A) by aligning Cαatoms of residues 353-371 of SARS-CoV spike-S230 structure (PDB 6NB6, chain E) and overlaying the glycan at N357_RBD_ from the SARS-CoV spike on the RBD model of the C118-RBD crystal structure. Sequence alignments were done using the MUSCLE server (https://www.ebi.ac.uk/Tools/msa/muscle/) (Edgar, 2004). Secondary structure was defined as described in (Huo et al., 2020).

To predict whether intra-spike crosslinking by a single IgG binding to a spike trimer might be possible, we measured the distance between residue 222_HC_ Cα atoms in the C_H_1 domains of adjacent Fabs in Fab-S structures as previously described (Barnes et al., 2020a). This distance was compared to analogous distances in crystal structures of intact IgGs (42Å, PDB 1HZH; 48Å, PDB 1IGY; 52Å, PDB 1IGT). We accounted for potential influences of crystal packing in intact IgG structures, flexibility in the V_H_-V_L_/C_H_1-C_L_ elbow bend angle, and uncertainties in C_H_1-C_L_ domain placement in Fab-S cryo-EM structures, by setting a cut-off of ≤65Å for this measured distance as potentially allowing for a single IgG to include both Fabs when binding a spike trimer.

## Supplemental Items

**Table S1:** Crystallographic data collection and validation statistics for C118-SARS RBD and C022-SARS2 RBD (related to Figure 2)

| **PDB ID** | **C118 Fab – SARS-CoV RBD**<br>**AAAA** | **C022 Fab – SARS-CoV-2 RBD**<br>**BBBB** |
| --- | --- | --- |
| **Data collection<sup>a,b</sup>** |  |  |
| Space group | P2<sub>1</sub> | P6<sub>1</sub> |
| Unit cell (Å) | 92.9, 90.0, 93.9 | 178.4 178.4 247.3 |
| α, β, γ (°) | 90.0, 113.3, 90.0 | 90.0 90.0 120.0 |
| Wavelength (Å) | 0.980 | 0.980 |
| Resolution (Å) | 46.6 -2.70 (2.80-2.70) | 44.6-3.20 (3.32-3.20) |
| Unique Reflections | 37504 (3085) | 73262 (7338) |
| Completeness (%) | 95.7 (79.5) | 99.5 (96.9) |
| Redundancy | 3.4 (2.9) | 10.3 (9.8) |
| CC<sub>1/2</sub> (%) | 99.8 (91.8) | 99.8 (27.7) |
| <I/σI> | 13.14 (2.27) | 12.87 (0.56) |
| R<sub>merge</sub> (%) | 6.22 (40.1) | 16.4 (492) |
| R<sub>pim</sub> (%) | 3.96 (28.0) | 5.36 (165) |
| Wilson B-factor (Å<sup>2</sup>) | 59.53 | 125.06 |
| **Refinement and Validation** |  |  |
| Resolution (Å) | 2.70 | 3.20 |
| Number of atoms | 9582 | 19802 |
| Protein | 9516 | 19696 |
| Ligand | 66 | 106 |
| Waters | 0 | 0 |
| R<sub>work</sub>/R<sub>free</sub> (%) | 19.2/24.1 | 18.8/23.1 |
| R.m.s. deviations |  |  |
| Bond lengths (Å) | 0.009 | 0.010 |
| Bond angles (°) | 1.16 | 1.23 |
| MolProbity score | 1.96 | 2.04 |
| Clashscore (all atom) | 7.61 | 8.67 |
| Poor rotamers (%) | 1.71 | 1.77 |
| Ramachandran plot |  |  |
| Favored (%) | 94.60 | 94.36 |
| Allowed (%) | 5.07 | 5.28 |
| Disallowed (%) | 0.33 | 0.36 |
| Average B-factor (Å<sup>2</sup>) | 63.16 | 124.1 |

<sup>a</sup> Data collected on 12-2 beamline at the Stanford Synchrotron Radiation Lightsource (SSRL).
<sup>b</sup> Numbers in parentheses correspond to the highest resolution shell.

**Table S2.** Buried Surface Area (related to Figure 3)

| **Interface Buried Surface Area (Å<sup>2</sup>)** |  |  |  |  |
| --- | --- | --- | --- | --- |
| **Structure**<br>**PDB** | **C118 Fab/**<br>**SARS RBD**<br>**this study** | **C022 Fab/**<br>**SARS2 RBD**<br>**this study** | **CR3022 Fab/**<br>**SARS2 RBD**<br>**PDB 6W41** | **COVA1-16 Fab/**<br>**SARS2 RBD**<br>**PDB 7JMW** |
| **Heavy Chain**<br>**Paratope** | 750 | 769 | 593 | 673 |
| **FWRH1** | 0 | 12 | 4 | 0 |
| **CDRH1** | 197 | 218 | 271 | 109 |
| **FWRH2** | 0 | 0 | 0 | 0 |
| **CDRH2** | 92 | 3 | 113 | 0 |
| **FWRH3** | 0 | 0 | 3 | 0 |
| **CDRH3** | 461 | 537 | 202 | 564 |
| **FWRH4** | 0 | 0 | 0 | 0 |
| **Light Chain**<br>**Paratope** | 272 | 200 | 432 | 154 |
| **FWRL1** | 0 | 0 | 0 | 0 |
| **CDRL1** | 0 | 1 | 239 | 0 |
| **FWRL2** | 4 | 48 | 50 | 34 |
| **CDRL2** | 121 | 20 | 62 | 0 |
| **FWRL3** | 147 | 130 | 28 | 120 |
| **CDRL3** | 0 | 0 | 0 | 0 |
| **FWRL4** | 0 | 0 | 0 | 0 |
| **Total Paratope** | 1022 | 969 | 1025 | 827 |
| **Heavy Chain**<br>**Epitope** | 656 | 704 | 593 | 628 |
| **Light Chain Epitope** | 292 | 202 | 398 | 153 |
| **Total Epitope** | 948 | 906 | 990 | 780 |

**Table S3.**
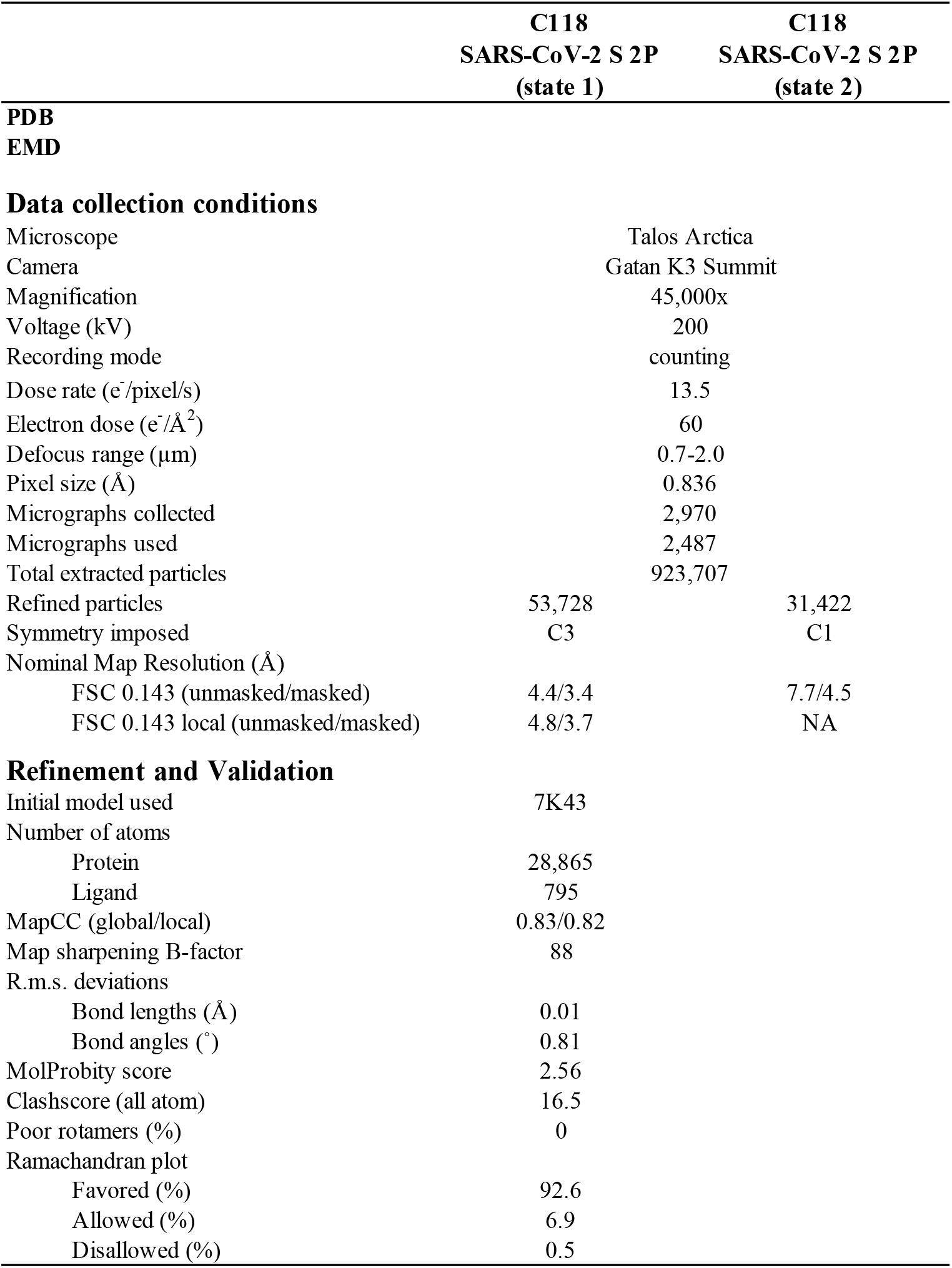
Cryo-EM data collection and refinement statistics for C118-S complex structure (related to Figure 4).

**Figure S1.**
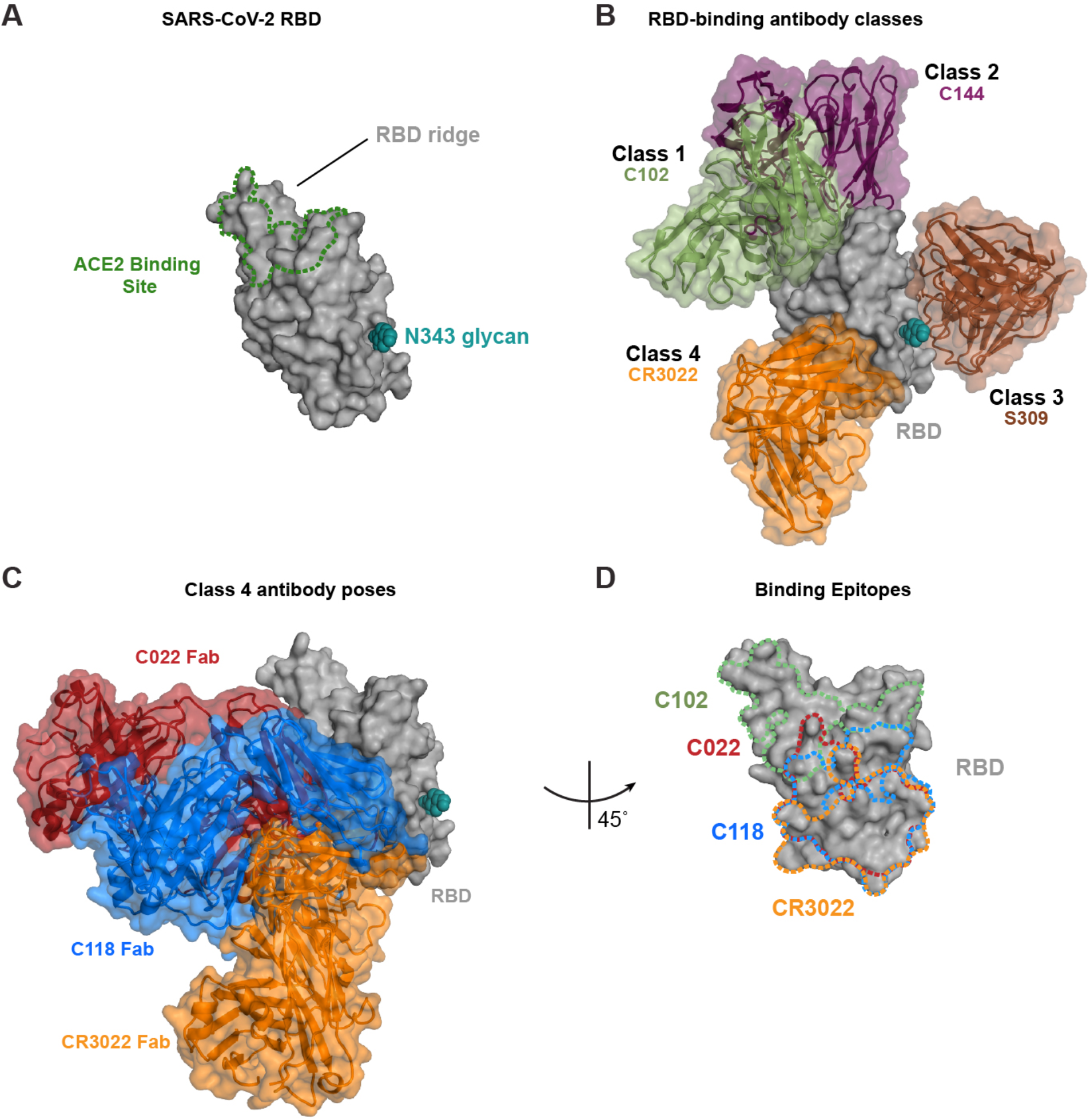
Epitopes of class 1 – class 4 anti-RBD antibodies (related to Figures 1 and 2) (A) SARS-CoV-2 RBD surface representation (grey) with N343 glycan (teal). The ACE2 binding site is represented by a green dashed line. (B) SARS-CoV-2 RBD surface representation (grey) with overlaid bound models of V_H_V_L_ of antibodies for class 1 (C102, green, PDB: 7K8M), class 2 (C144, purple, PDB: 7K90), Class 3 (S309, brown, 7JMX), and class 4 (CR3022, orange, PDB: 6W41). (C) SARS-CoV-2 surface representation with overlay of bound C118 Fab (blue), C022 Fab (red), and CR3022 Fab (orange, PDB: 6W41). (D) SARS-CoV-2 RBD surface representation (grey) at 45^°^ angle from previous panels. Dotted outlines show epitopes of C102 (green), C022 (red), C118 (blue), and CR3022 (orange) mapped onto SARS-CoV-2 RBD.

**Figure S2.**
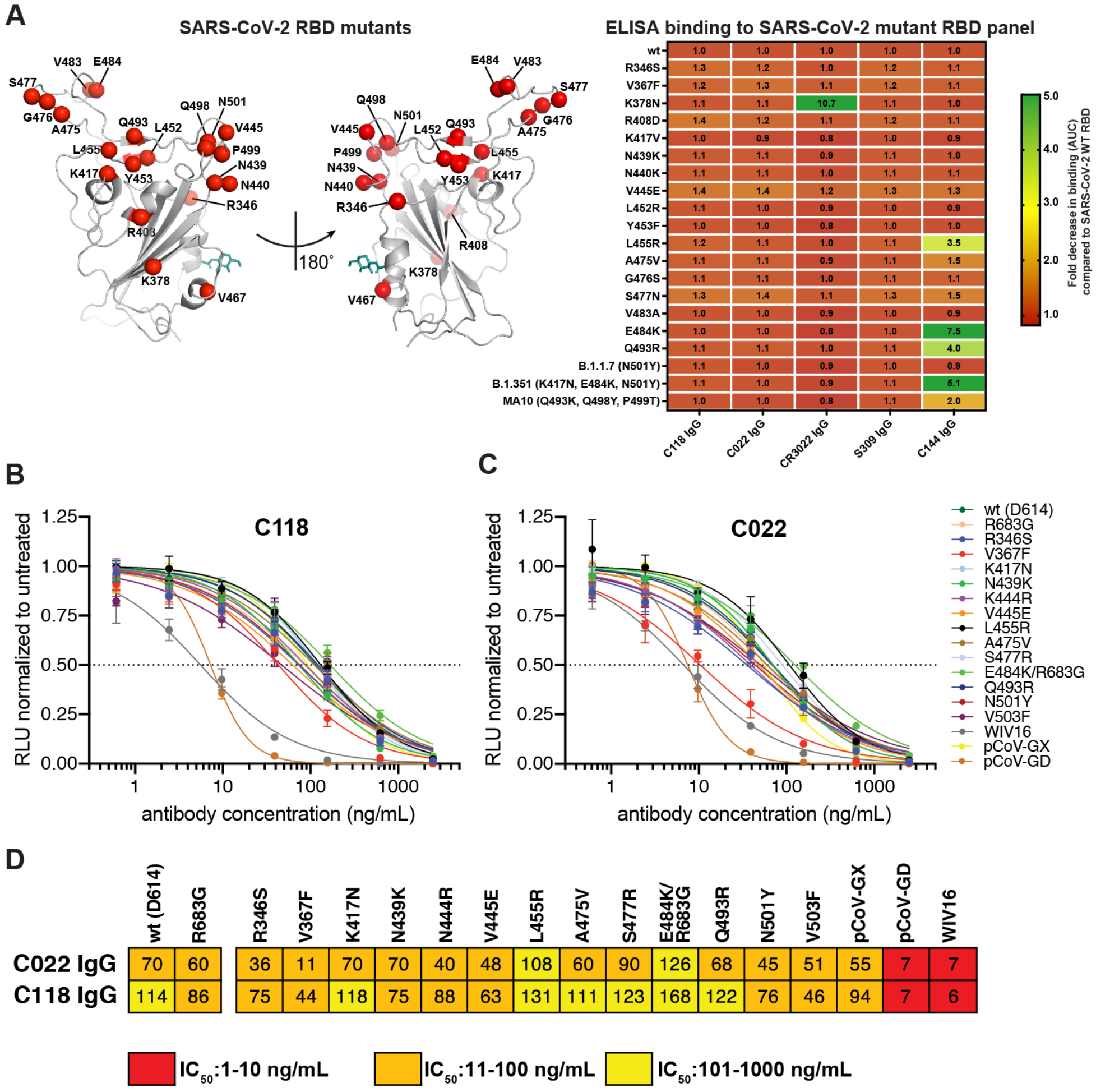
Binding and Neutralization of SARS-CoV-2 RBD or pseudoviruses carrying spike amino acid substitutions as well as other sarbecovirus pseudoviruses by C118 and C022 (related to Figure 1). (A) Left: Cartoon model of SARS-CoV-2 RBD (RBD from C022-RBD structure) showing locations of point mutations as red spheres and the N343_RBD_ *N*-glycan as teal sticks. Right: Comparison of binding of the indicated monoclonal IgGs to RBD mutants from ELISA data shown as AUC values normalized to antibody binding to ‘wt’ SARS-CoV-2 RBD. Data presented are normalized mean AUC values from two independent experiments. (B-C) Normalized relative luminescence values for cell lysates of HT1080_ACE2_ cells 48h after infection with SARS-CoV-2 pseudovirus carrying indicated spike variants in the presence of increasing concentrations of monoclonal IgGs C118 (B) and C022 (C). The mutants represented substitutions found in circulating SARS-CoV-2 sequences with frequencies >0.01% in GISAID (Shu and McCauley, 2017). Mean and standard deviation of two experiments, each performed in duplicate (n=4), is shown. (D) Half-maximal inhibitory concentrations (IC_50_) calculated from the neutralization curves in panels B and C for monoclonal IgGs C022 and C118 for neutralization of ‘wt’ (D614 S trimer) and the indicated mutant SARS-CoV-2 S pseudotyped viruses, as well as other sarbecovirus pseudoviruses. IC_50_ values are means of 2 independent experiments. Colors indicate IC_50_ ranges, as indicated. The E484K substitution was constructed in an R683G (furin cleavage site mutant) background to increase infectivity (Muecksch et al., 2021).

**Figure S3.**
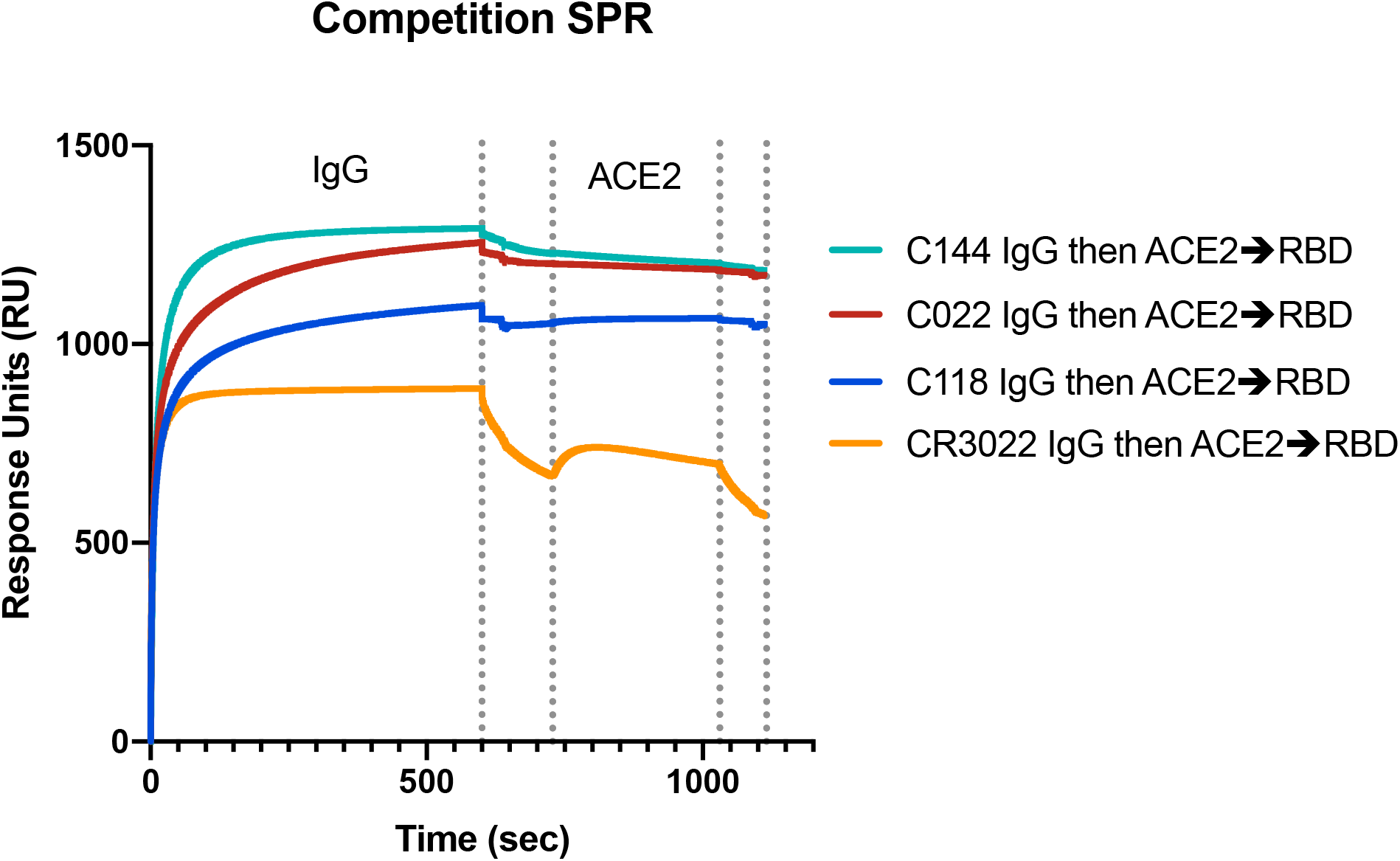
SPR-based competition experiment. SARS-CoV-2 RBD was coupled to a biosensor chip using primary amine chemistry. An IgG was injected first (seconds 0-600). Seconds 600 - 730 represent the delay required to switch samples for a subsequent injection. From seconds 730 – 1030, soluble ACE2 was injected. Buffer was injected after 1030 seconds.

**Figure S4.**
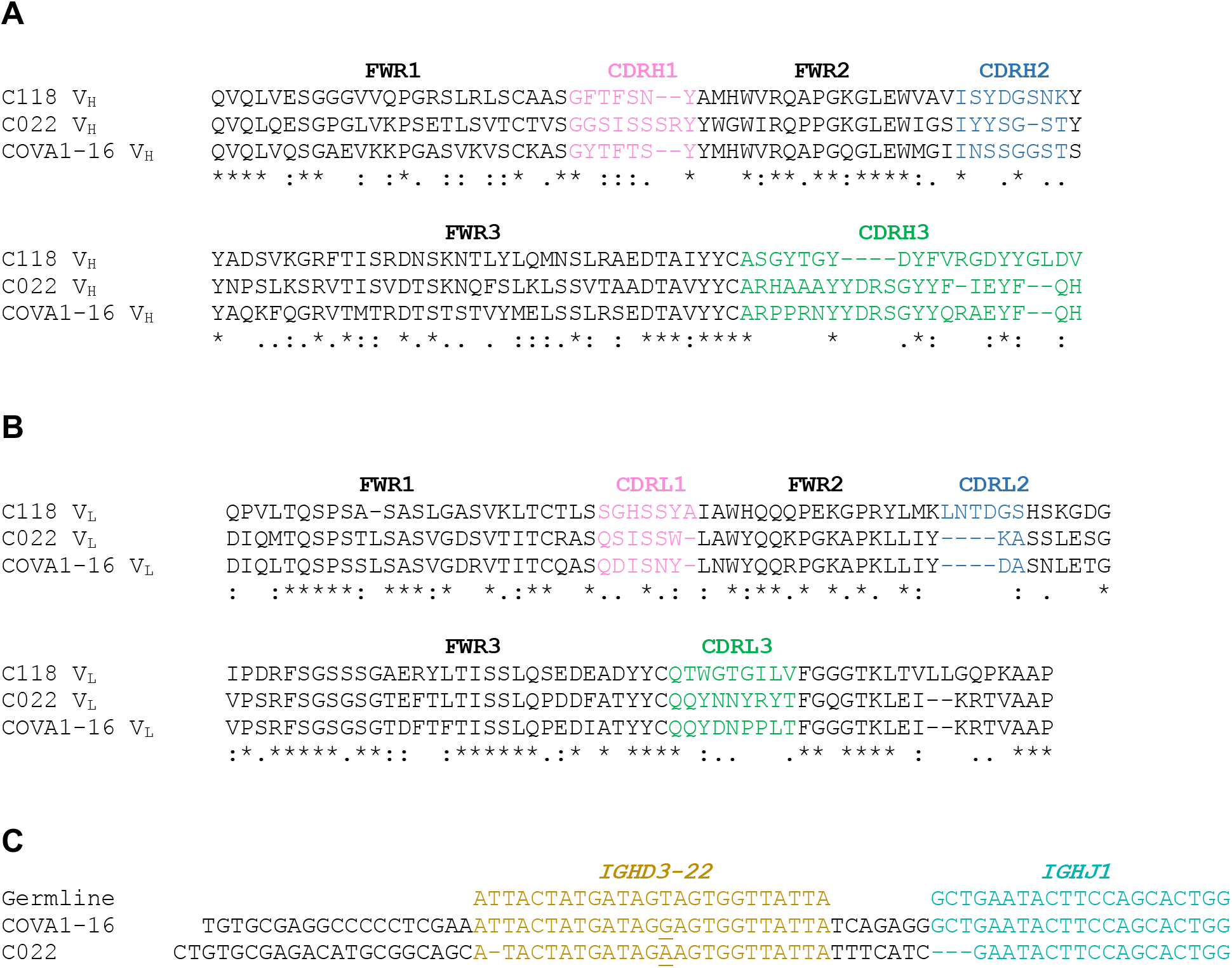
Comparison of C022, C118, and COVA1-16 V_H_V_L_ sequences (related to Figure 3) (A,B) Alignment of C118, C022, and COVA1-16 (A) V_H_ and (B) V_L_ domains. Framework regions (FWRs) and complementarity determining regions (CDRs) assigned using the IMGT definition (Lefranc et al., 2015). Conserved residues (*), residues with similar properties (:), residues with weakly similar properties (.). (C) Alignment of portions of the C022 and COVA1-16 V_H_ domain nucleotide sequences to germline gene segment sequences for *IGHD3-22* (sand) and *IGHJ1* (teal).

**Figure S5.**
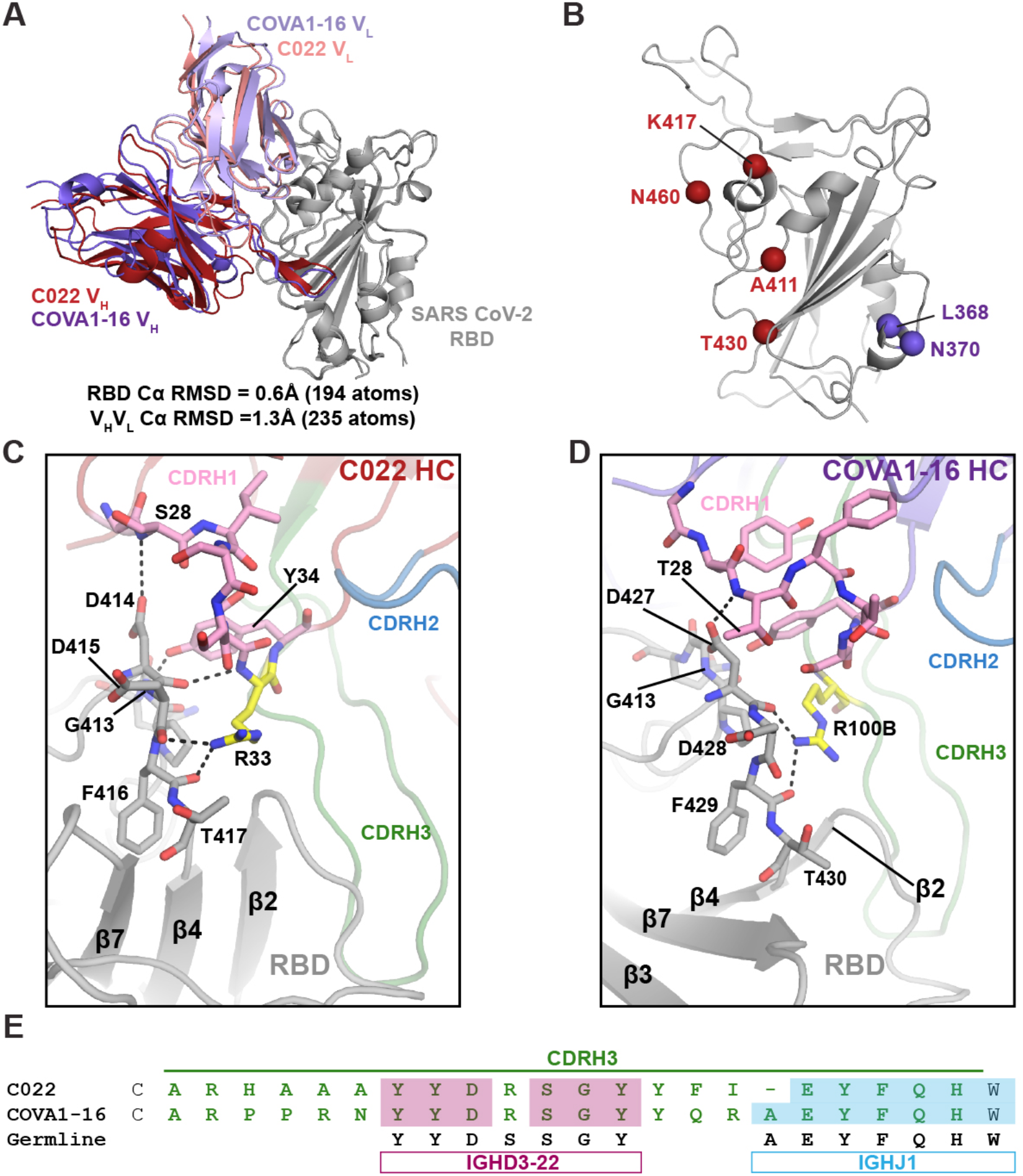
Comparison of C022 and COVA1-16 antibody binding. (A) Cartoon representation of superposition of RBDs from C022-RBD and COVA1-16-RBD (PDB 7JMW) crystal structures aligned by their Cα atoms. (B) SARS-CoV-2 RBD cartoon model showing differences between C022 (red) and COVA1-16 (purple) epitopes. (C,D) Interacting residues of SARS-CoV-2 RBD with the CDRH1 regions from (D) C022 Fab and (E) COVA1-16 Fab (PDB 7JMW). Colors: Oxygens (red), nitrogens (blue). Carbon atoms of critical arginines from each paratope are highlighted in yellow. (E) Alignment of CDRH3 amino acid sequences for C022 and COVA1-16 and germline-encoded amino acids derived from *IGD3-22* (mauve) and *IGJ1* (blue) gene segments. Identities between C022 or COVA1-16 with *IGHD3-22* and *IGHJ1* amino acid sequences shown as shaded boxes and residues within the CDRH3 loop are green.

**Figure S6.**
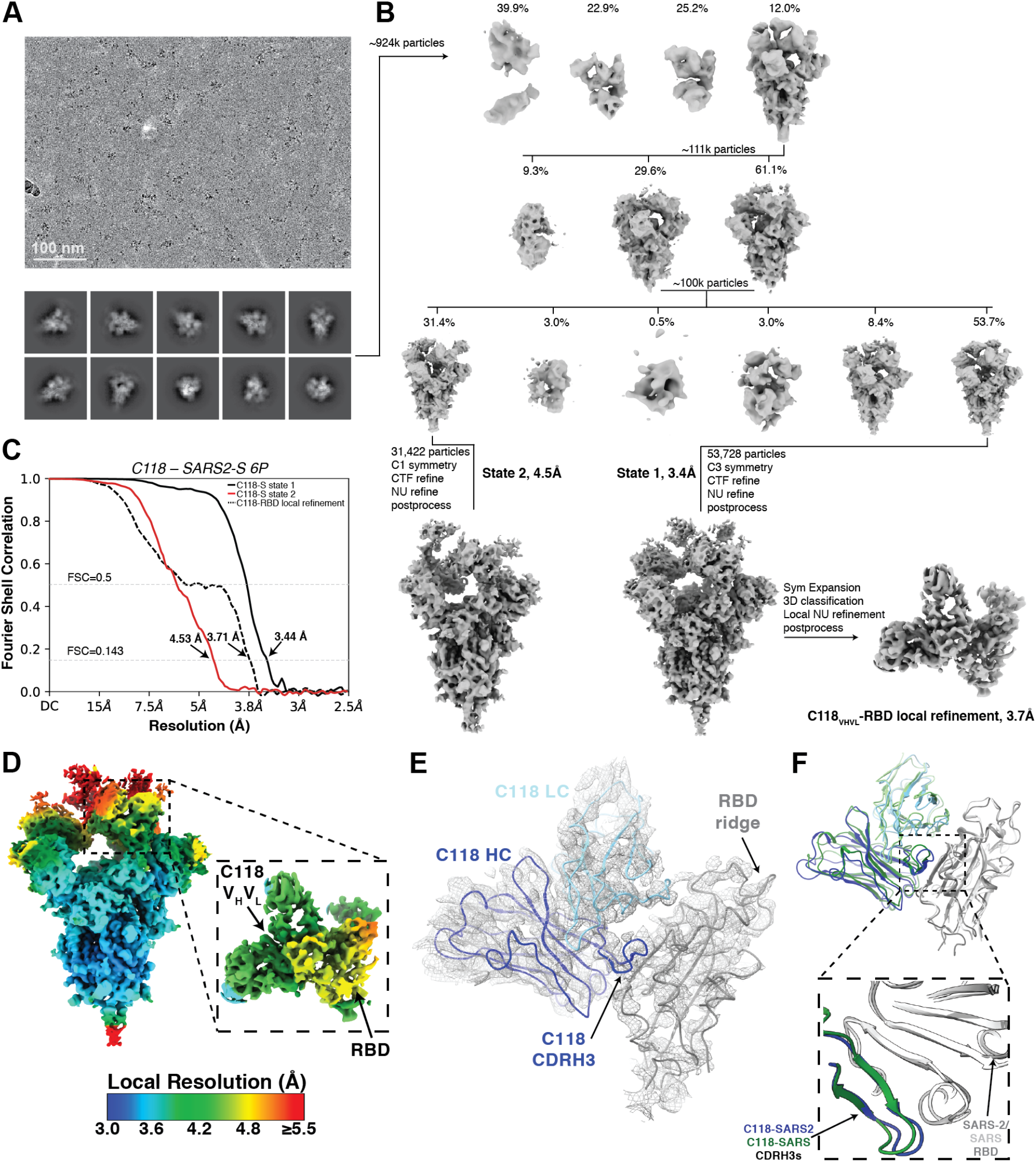
Details for cryo-EM C118-S reconstructions (related to Figure 5) (A) C118-S 6P representative micrograph (top) and 2D class averages (bottom). (B) Data processing pipeline for C118-S structure and focused refinement of C118-RBD portion of the structure. (C) Gold Standard FSC for final reconstruction of C118-S reconstructions. Resolutions at FSC=0.143 are shown for each volume. (D) Local resolution estimates for C118-S map and C118-RBD focused map calculated in cryoSPARC. (E) Ribbon representation of C118-RBD rigid body-refined model into cryoEM density contoured at 8σ (gray mesh). (F) Overlay of C118-RBD (SARS-CoV-2) from Fab-S cryo-EM structure (blue and gray cartoon) and C118-RBD (SARS-CoV) (green and light gray). CDRH3 interactions are highlighted.

## References

Adams, P.D., Afonine, P.V., Bunkoczi, G., Chen, V.B., Davis, I.W., Echols, N., Headd, J.J., Hung, L.W., Kapral, G.J., Grosse-Kunstleve, R.W., et al. (2010). PHENIX: a comprehensive Python-based system for macromolecular structure solution. Acta Crystallogr D Biol Crystallogr 66, 213–221.

Annavajhala, M.K., Mohri, H., Zucker, J.E., Sheng, Z., Wang, P., Gomez-Simmonds, A., Ho, D.D., and Uhlemann, A.-C. (2021). A Novel SARS-CoV-2 Variant of Concern, B.1.526, Identified in New York. medRxiv 10.1101/2021.02.23.21252259.

Barnes, C.O., Jette, C.A., Abernathy, M.E., Dam, K.-M.A., Esswein, S.R., Gristick, H.B., Malyutin, A.G., Sharaf, N.G., Huey-Tubman, K.E., Lee, Y.E., et al. (2020a). SARS-CoV-2 neutralizing antibody structures inform therapeutic strategies. Nature 588, 682–687.

Barnes, C.O., West, A.P., Jr., Huey-Tubman, K.E., Hoffmann, M.A.G., Sharaf, N.G., Hoffman, P.R., Koranda, N., Gristick, H.B., Gaebler, C., Muecksch, F., et al. (2020b). Structures of Human Antibodies Bound to SARS-CoV-2 Spike Reveal Common Epitopes and Recurrent Features of Antibodies. Cell 182, 828–842 e816.

Bell, J.M., Chen, M., Baldwin, P.R., and Ludtke, S.J. (2016). High resolution single particle refinement in EMAN2.1. Methods 100, 25–34.

Brouwer, P.J.M., Caniels, T.G., van der Straten, K., Snitselaar, J.L., Aldon, Y., Bangaru, S., Torres, J.L., Okba, N.M.A., Claireaux, M., Kerster, G., et al. (2020). Potent neutralizing antibodies from COVID-19 patients define multiple targets of vulnerability. Science 369, 643–650.

Bunkóczi, G., and Read, R.J. (2011). Improvement of molecular-replacement models with Sculptor. Acta Cryst 67, 303–312.

Cao, Y., Su, B., Guo, X., Sun, W., Deng, Y., Bao, L., Zhu, Q., Zhang, X., Zheng, Y., Geng, C., et al. (2020). Potent neutralizing antibodies against SARS-CoV-2 identified by high-throughput single-cell sequencing of convalescent patients’ B cells. Cell 10.1016/j.cell.2020.05.025.

Chan, K.K., Dorosky, D., Sharma, P., Abbasi, S.A., Dye, J.M., Kranz, D.M., Herbert, A.S., and Procko, E. (2020). Engineering human ACE2 to optimize binding to the spike protein of SARS coronavirus 2. Science 369, 1261–1265.

Chen, V.B., Arendall, W.B., 3rd, Headd, J.J., Keedy, D.A., Immormino, R.M., Kapral, G.J., Murray, L.W., Richardson, J.S., and Richardson, D.C. (2010). MolProbity: all-atom structure validation for macromolecular crystallography. Acta Crystallogr D Biol Crystallogr 66, 12–21.

Cohen, A.A., Gnanapragasam, P.N.P., Lee, Y.E., Hoffman, P.R., Ou, S., Kakutani, L.M., Keeffe, J.R., Wu, H.-J., Howarth, M., West, A.P., et al. (2021). Mosaic nanoparticles elicit cross-reactive immune responses to zoonotic coronaviruses in mice. Science 371, 735–741.

Corti, D., Suguitan, A.L., Pinna, D., Silacci, C., Fernandez-Rodriguez, B.M., Vanzetta, F., Santos, C., Luke, C.J., Torres-Velez, F.J., Temperton, N.J., et al. (2010). Heterosubtypic neutralizing antibodies are produced by individuals immunized with a seasonal influenza vaccine. Journal of Clinical Investigation 120, 1663–1673.

Crawford, K.H.D., Eguia, R., Dingens, A.S., Loes, A.N., Malone, K.D., Wolf, C.R., Chu, H.Y., Tortorici, M.A., Veesler, D., Murphy, M., et al. (2020). Protocol and Reagents for Pseudotyping Lentiviral Particles with SARS-CoV-2 Spike Protein for Neutralization Assays. Viruses 12.

de Wit, E., van Doremalen, N., Falzarano, D., and Munster, V.J. (2016). SARS and MERS: recent insights into emerging coronaviruses. Nat Rev Microbiol 14, 523–534.

Dejnirattisai, W., Zhou, D., Ginn, H.M., Duyvesteyn, H.M.E., Supasa, P., Case, J.B., Zhao, Y., Walter, T.S., Mentzer, A.J., Liu, C., et al. (2021). The antigenic anatomy of SARS-CoV-2 receptor binding domain. Cell 10.1016/j.cell.2021.02.032.

Desrosiers, R.C., Wagh, K., Bhattacharya, T., Williamson, C., Robles, A., Bayne, M., Garrity, J., Rist, M., Rademeyer, C., Yoon, H., et al. (2016). Optimal Combinations of Broadly Neutralizing Antibodies for Prevention and Treatment of HIV-1 Clade C Infection. PLOS Pathogens 12, e1005520.

Edgar, R.C. (2004). MUSCLE: multiple sequence alignment with high accuracy and high throughput. Nuc Acids Res 32, 1792–1797.

Emsley, P., Lohkamp, B., Scott, W.G., and Cowtan, K. (2010). Features and development of Coot. Acta Crystallogr D Biol Crystallogr 66, 486–501.

Faria, N.R., Claro, I.M., Candido, D., Moyses Franco, L.A., Andrade, P.S., Coletti, T.M., Silva, C.A.M., Sales, F.C., Manuli, E.R., Agular, R.S., et al. (2021). Genomic characterisation of an emergent SARS-CoV-2 lineage in Manaus: preliminary findings. https://virological.org/t/genomic-characterisation-of-an-emergent-sars-cov-2-lineage-in-manaus-preliminary-findings/586.

Fung, T.S., and Liu, D.X. (2019). Human Coronavirus: Host-Pathogen Interaction. Annu Rev Microbiol 73, 529–557.

Goddard, T.D., Huang, C.C., and Ferrin, T.E. (2007). Visualizing density maps with UCSF Chimera. J Struct Biol 157, 281–287.

Goddard, T.D., Huang, C.C., Meng, E.C., Pettersen, E.F., Couch, G.S., Morris, J.H., and Ferrin, T.E. (2018). UCSF ChimeraX: Meeting modern challenges in visualization and analysis. Protein Sci 27, 14–25.

Greaney, A.J., Loes, A.N., Crawford, K.H.D., Starr, T.N., Malone, K.D., Chu, H.Y., and Bloom, J.D. (2021). Comprehensive mapping of mutations in the SARS-CoV-2 receptor-binding domain that affect recognition by polyclonal human plasma antibodies. Cell Host & Microbe 29, 463–476.e466.

Gristick, H.B., von Boehmer, L., West, A.P., Jr., Schamber, M., Gazumyan, A., Golijanin, J., Seaman, M.S., Fatkenheuer, G., Klein, F., Nussenzweig, M.C., et al. (2016). Natively glycosylated HIV-1 Env structure reveals new mode for antibody recognition of the CD4-binding site. Nat Struct Mol Biol 23, 906–915.

Guindon, S., Dufayard, J.F., Lefort, V., Anisimova, M., Hordijk, W., and Gascuel, O. (2010). New algorithms and methods to estimate maximum-likelihood phylogenies: assessing the performance of PhyML 3.0. Syst Biol 59, 307–321.

Haagmans, B.L., Al Dhahiry, S.H.S., Reusken, C.B.E.M., Raj, V.S., Galiano, M., Myers, R., Godeke, G.-J., Jonges, M., Farag, E., Diab, A., et al. (2014). Middle East respiratory syndrome coronavirus in dromedary camels: an outbreak investigation. The Lancet Infectious Diseases 14, 140–145.

Hoffmann, M., Arora, P., Groß, R., Seidel, A., Hörnich, B.F., Hahn, A.S., Krüger, N., Graichen, L., Hofmann-Winkler, H., Kempf, A., et al. (2021). SARS-CoV-2 variants B.1.351 and P.1 escape from neutralizing antibodies. Cell 10.1016/j.cell.2021.03.036.

Hoffmann, M., Kleine-Weber, H., Schroeder, S., Kruger, N., Herrler, T., Erichsen, S., Schiergens, T.S., Herrler, G., Wu, N.H., Nitsche, A., et al. (2020). SARS-CoV-2 Cell Entry Depends on ACE2 and TMPRSS2 and Is Blocked by a Clinically Proven Protease Inhibitor. Cell 181, 271–280 e278.

Hsieh, C.-L., Goldsmith, J.A., Schaub, J.M., DiVenere, A.M., Kuo, H.-C., Javanmardi, K., Le, K.C., Wrapp, D., Lee, A.G., Liu, Y., et al. (2020). Structure-based design of prefusion-stabilized SARS-CoV-2 spikes. Science 369, 1501–1505.

Huo, J., Zhao, Y., Ren, J., Zhou, D., Duyvesteyn, H.M.E., Ginn, H.M., Carrique, L., Malinauskas, T., Ruza, R.R., Shah, P.N.M., et al. (2020). Neutralization of SARS-CoV-2 by Destruction of the Prefusion Spike. Cell Host & Microbe 28, 445–454.e446.

Kabsch, W. (2010). XDS. Acta Crystallogr D Biol Crystallogr 66, 125–132.

Kirchdoerfer, R.N., Cottrell, C.A., Wang, N., Pallesen, J., Yassine, H.M., Turner, H.L., Corbett, K.S., Graham, B.S., McLellan, J.S., and Ward, A.B. (2016). Pre-fusion structure of a human coronavirus spike protein. Nature 531, 118–121.

Klein, J.S. (2009). Investigations in the design and characterization of HIV-1 neutralizing molecules (Pasadena: California Institute of Technology), pp. 166.

Klein, J.S., and Bjorkman, P.J. (2010). Few and far between: how HIV may be evading antibody avidity. PLoS Pathog 6, e1000908.

Korber, B., Fischer, W.M., Gnanakaran, S., Yoon, H., Theiler, J., Abfalterer, W., Hengartner, N., Giorgi, E.E., Bhattacharya, T., Foley, B., et al. (2020). Tracking Changes in SARS-CoV-2 Spike: Evidence that D614G Increases Infectivity of the COVID-19 Virus. Cell 182, 812–827.e819.

Kreer, C., Zehner, M., Weber, T., Ercanoglu, M.S., Gieselmann, L., Rohde, C., Halwe, S., Korenkov, M., Schommers, P., Vanshylla, K., et al. (2020). Longitudinal Isolation of Potent Near-Germline SARS-CoV-2-Neutralizing Antibodies from COVID-19 Patients. Cell 10.1016/j.cell.2020.06.044.

Krissinel, E., and Henrick, K. (2007). Inference of macromolecular assemblies from crystalline state. J Mol Biol 372, 774–797.

Lan, J., Ge, J., Yu, J., Shan, S., Zhou, H., Fan, S., Zhang, Q., Shi, X., Wang, Q., Zhang, L., et al. (2020). Structure of the SARS-CoV-2 spike receptor-binding domain bound to the ACE2 receptor. Nature 581, 215–220.

Landau, M., Mayrose, I., Rosenberg, Y., Glaser, F., Martz, E., Pupko, T., and Ben-Tal, N. (2005). ConSurf 2005: the projection of evolutionary conservation scores of residues on protein structures. Nucleic Acids Res 33, W299–302.

Lee, B., Huang, K.-Y.A., Tan, T.K., Chen, T.-H., Huang, C.-G., Harvey, R., Hussain, S., Chen, C.-P., Harding, A., Gilbert-Jaramillo, J., et al. (2021). Breadth and function of antibody response to acute SARS-CoV-2 infection in humans. PLOS Pathogens 17, e1009352.

Lefranc, M.P., Giudicelli, V., Duroux, P., Jabado-Michaloud, J., Folch, G., Aouinti, S., Carillon, E., Duvergey, H., Houles, A., Paysan-Lafosse, T., et al. (2015). IMGT(R), the international ImMunoGeneTics information system(R) 25 years on. Nucleic Acids Res 43, D413–422.

Lefranc, M.P., Giudicelli, V., Ginestoux, C., Jabado-Michaloud, J., Folch, G., Bellahcene, F., Wu, Y., Gemrot, E., Brochet, X., Lane, J., et al. (2009). IMGT, the international ImMunoGeneTics information system. Nucleic Acids Res 37, D1006–1012.

Leist, S.R., Dinnon, K.H., Schäfer, A., Tse, L.V., Okuda, K., Hou, Y.J., West, A., Edwards, C.E., Sanders, W., Fritch, E.J., et al. (2020). A Mouse-Adapted SARS-CoV-2 Induces Acute Lung Injury and Mortality in Standard Laboratory Mice. Cell 183, 1070–1085.e1012.

Li, Q., Wu, J., Nie, J., Zhang, L., Hao, H., Liu, S., Zhao, C., Zhang, Q., Liu, H., Nie, L., et al. (2020). The Impact of Mutations in SARS-CoV-2 Spike on Viral Infectivity and Antigenicity. Cell 182, 1284–1294.e1289.

Li, W., Moore, M.J., Vasilieva, N., Sui, J., Wong, S.K., Berne, M.A., Somasundaran, M., Sullivan, J.L., Luzuriaga, K., Greenough, T.C., et al. (2003). Angiotensin-converting enzyme 2 is a functional receptor for the SARS coronavirus. Nature 426, 450–454.

Li, W., Shi, Z., Yu, M., Ren, W., Smith, C., Epstein, J.H., Wang, H., Crameri, G., Hu, Z., Zhang, H., et al. (2005). Bats are natural reservoirs of SARS-like coronaviruses. Science 310, 676–679.

Li, Z., Tomlinson, A.C., Wong, A.H., Zhou, D., Desforges, M., Talbot, P.J., Benlekbir, S., Rubinstein, J.L., and Rini, J.M. (2019). The human coronavirus HCoV-229E S-protein structure and receptor binding. Elife 8.

Liu, H., Wu, N.C., Yuan, M., Bangaru, S., Torres, J.L., Caniels, T.G., van Schooten, J., Zhu, X., Lee, C.-C.D., Brouwer, P.J.M., et al. (2020a). Cross-Neutralization of a SARS-CoV-2 Antibody to a Functionally Conserved Site Is Mediated by Avidity. Immunity 53, 1272–1280.e1275.

Liu, H., Yuan, M., Huang, D., Bangaru, S., Zhao, F., Lee, C.-C.D., Peng, L., Barman, S., Zhu, X., Nemazee, D., et al. (2021). A combination of cross-neutralizing antibodies synergizes to prevent SARS-CoV-2 and SARS-CoV pseudovirus infection. Cell Host & Microbe 10.1016/j.chom.2021.04.005.

Liu, L., Wang, P., Nair, M.S., Yu, J., Rapp, M., Wang, Q., Luo, Y., Chan, J.F.W., Sahi, V., Figueroa, A., et al. (2020b). Potent neutralizing antibodies against multiple epitopes on SARS-CoV-2 spike. Nature 584, 450–456.

Mastronarde, D.N. (2005). Automated electron microscope tomography using robust prediction of specimen movements. J Struct Biol 152, 36–51.

McCoy, A.J., Grosse-Kunstleve, R.W., Adams, P.D., Winn, M.D., Storoni, L.C., and Read, R.J. (2007). Phaser crystallographic software. J Appl Crystallogr 40, 658–674.

Muecksch, F., Weisblum, Y., Barnes, C.O., Schmidt, F., Schaefer-Babajew, D., Lorenzi, J.C.C., Flyak, A.I., DeLaitsch, A.T., Huey-Tubman, K.E., Hou, S., et al. (2021). Development of potency, breadth and resilience to viral escape mutations in SARS-CoV-2 neutralizing antibodies. bioRxiv 10.1101/2021.03.07.434227.

Piccoli, L., Park, Y.-J., Tortorici, M.A., Czudnochowski, N., Walls, A.C., Beltramello, M., Silacci-Fregni, C., Pinto, D., Rosen, L.E., Bowen, J.E., et al. (2020a). Mapping Neutralizing and Immunodominant Sites on the SARS-CoV-2 Spike Receptor-Binding Domain by Structure-Guided High-Resolution Serology. Cell 183, 1024–1042.e1021.

Piccoli, L., Park, Y.-J., Tortorici, M.A., Czudnochowski, N., Walls, A.C., Beltramello, M., Silacci-Fregni, C., Pinto, D., Rosen, L.E., Bowen, J.E., et al. (2020b). Mapping neutralizing and immunodominant sites on the SARS-CoV-2 spike receptor-binding domain by structure-guided high-resolution serology. Cell 10.1016/j.cell.2020.09.037.

Pinto, D., Park, Y.-J., Beltramello, M., Walls, A.C., Tortorici, M.A., Bianchi, S., Jaconi, S., Culap, K., Zatta, F., De Marco, A., et al. (2020). Cross-neutralization of SARS-CoV-2 by a human monoclonal SARS-CoV antibody. Nature 583, 290–295.

Punjani, A., Rubinstein, J.L., Fleet, D.J., and Brubaker, M.A. (2017). cryoSPARC: algorithms for rapid unsupervised cryo-EM structure determination. Nat Methods 14, 290–296.

Rambaut, A., Pybus, O., Barclay, W., Barrett, J., Carabelli, A., Connor, T., Peacock, T., Robertson, D.L., Volz, E., and UK, C.-G.C. (2020). Preliminary genomic characterisation of an emergent SARS-CoV-2 lineage in the UK defined by a novel set of spike mutations. virologicalorg, https://virological.org/t/preliminary-genomic-characterisation-of-an-emergent-sars-cov-2-lineage-in-the-uk-defined-by-a-novel-set-of-spike-mutations/563.

Rappazzo, C.G., Tse, L.V., Kaku, C.I., Wrapp, D., Sakharkar, M., Huang, D., Deveau, L.M., Yockachonis, T.J., Herbert, A.S., Battles, M.B., et al. (2021). Broad and potent activity against SARS-like viruses by an engineered human monoclonal antibody. Science 371, 823–829.

Robbiani, D.F., Gaebler, C., Muecksch, F., Lorenzi, J.C.C., Wang, Z., Cho, A., Agudelo, M., Barnes, C.O., Gazumyan, A., Finkin, S., et al. (2020). Convergent antibody responses to SARS-CoV-2 in convalescent individuals. Nature 584, 437–442.

Rogers, T.F., Zhao, F., Huang, D., Beutler, N., Burns, A., He, W.T., Limbo, O., Smith, C., Song, G., Woehl, J., et al. (2020). Rapid isolation of potent SARS-CoV-2 neutralizing antibodies and protection in a small animal model. Science 10.1126/science.abc7520.

Sauer, M.M., Tortorici, M.A., Park, Y.-J., Walls, A.C., Homad, L., Acton, O., Bowen, J., Wang, C., Xiong, X., de van der Schueren, W., et al. (2021). Structural basis for broad coronavirus neutralization. bioRxiv 10.1101/2020.12.29.424482.

Schaefer, W., Regula, J.T., Bahner, M., Schanzer, J., Croasdale, R., Durr, H., Gassner, C., Georges, G., Kettenberger, H., Imhof-Jung, S., et al. (2011). Immunoglobulin domain crossover as a generic approach for the production of bispecific IgG antibodies. Proc Natl Acad Sci U S A 108, 11187–11192.

Scharf, L., Wang, H., Gao, H., Chen, S., McDowall, A.W., and Bjorkman, P.J. (2015). Broadly Neutralizing Antibody 8ANC195 Recognizes Closed and Open States of HIV-1 Env. Cell 162, 1379–1390.

Scheid, J.F., Mouquet, H., Ueberheide, B., Diskin, R., Klein, F., Olivera, T.Y., Pietzsch, J., Fenyo, D., Abadir, A., Velinzon, K., et al. (2011). Sequence and Structural Convergence of Broad and Potent HIV Antibodies That Mimic CD4 Binding. Science 333, 1633–1637.

Schmidt, F., Weisblum, Y., Muecksch, F., Hoffmann, H.-H., Michailidis, E., Lorenzi, J.C.C., Mendoza, P., Rutkowska, M., Bednarski, E., Gaebler, C., et al. (2020). Measuring SARS-CoV-2 neutralizing antibody activity using pseudotyped and chimeric viruses. Journal of Experimental Medicine 217.

Schoofs, T., Barnes, C.O., Suh-Toma, N., Golijanin, J., Schommers, P., Gruell, H., West, A.P., Jr., Bach, F., Lee, Y.E., Nogueira, L., et al. (2019). Broad and Potent Neutralizing Antibodies Recognize the Silent Face of the HIV Envelope. Immunity 50, 1513–1529 e1519.

Schrödinger, L. (2011). The PyMOL Molecular Graphics System (The PyMOL Molecular Graphics System).

Seydoux, E., Homad, L.J., MacCamy, A.J., Parks, K.R., Hurlburt, N.K., Jennewein, M.F., Akins, N.R., Stuart, A.B., Wan, Y.H., Feng, J., et al. (2020). Analysis of a SARS-CoV-2-Infected Individual Reveals Development of Potent Neutralizing Antibodies with Limited Somatic Mutation. Immunity 53, 98–105 e105.

Shah, P., Canziani, G.A., Carter, E.P., and Chaiken, I. (2021). The Case for S2: The Potential Benefits of the S2 Subunit of the SARS-CoV-2 Spike Protein as an Immunogen in Fighting the COVID-19 Pandemic. Frontiers in Immunology 12.

Shi, R., Shan, C., Duan, X., Chen, Z., Liu, P., Song, J., Song, T., Bi, X., Han, C., Wu, L., et al. (2020). A human neutralizing antibody targets the receptor-binding site of SARS-CoV-2. Nature 584, 120–124.

Shu, Y., and McCauley, J. (2017). GISAID: Global initiative on sharing all influenza data - from vision to reality. Euro Surveill 22.

Sievers, F., Wilm, A., Dineen, D., Gibson, T.J., Karplus, K., Li, W., Lopez, R., McWilliam, H., Remmert, M., Soding, J., et al. (2011). Fast, scalable generation of high-quality protein multiple sequence alignments using Clustal Omega. Mol Syst Biol 7, 539.

Song, H.D., Tu, C.C., Zhang, G.W., Wang, S.Y., Zheng, K., Lei, L.C., Chen, Q.X., Gao, Y.W., Zhou, H.Q., Xiang, H., et al. (2005). Cross-host evolution of severe acute respiratory syndrome coronavirus in palm civet and human. Proceedings of the National Academy of Sciences 102, 2430–2435.

Starr, T.N., Czudnochowski, N., Zatta, F., Park, Y.-J., Liu, Z., Addetia, A., Pinto, D., Beltramello, M., Hernandez, P., Greaney, A.J., et al. (2021a). Antibodies to the SARS-CoV-2 receptor-binding domain that maximize breadth and resistance to viral escape. bioRxiv 10.1101/2021.04.06.438709.

Starr, T.N., Greaney, A.J., Addetia, A., Hannon, W.W., Choudhary, M.C., Dingens, A.S., Li, J.Z., and Bloom, J.D. (2021b). Prospective mapping of viral mutations that escape antibodies used to treat COVID-19. Science 371, 850–854.

Tegally, H., Wilkinson, E., Giovanetti, M., Iranzadeh, A., Fonseca, V., Giandhari, J., Doolabh, D., Pillay, S., San, E.J., Msomi, N., et al. (2020). Emergence and rapid spread of a new severe acute respiratory syndrome-related coronavirus 2 (SARS-CoV-2) lineage with multiple spike mutations in South Africa. medRxiv 10.1101/2020.12.21.20248640.

Terwilliger, T.C., Adams, P.D., Afonine, P.V., and Sobolev, O.V. (2018). A fully automatic method yielding initial models from high-resolution cryo-electron microscopy maps. Nat Methods 15, 905–908.

Tortorici, M.A. (2020). Ultrapotent human antibodies protect against SARS-CoV-2 challenge via multiple mechanisms. Science 370, 950–957.

Tortorici, M.A., Czudnochowski, N., Starr, T.N., Marzi, R., Walls, A.C., Zatta, F., Bowen, J.E., Jaconi, S., iulio, J.d., Wang, Z., et al. (2021). 10.1101/2021.04.07.438818.

Voloch, C.M., Silva F, R.d., de Almeida, L.G.P., Cardoso, C.C., Brustolini, O.J., Gerber, A.L., Guimarães, A.P.d.C., Mariani, D., Costa, R.M.d., Ferreira, O.C., et al. (2020). Genomic characterization of a novel SARS-CoV-2 lineage from Rio de Janeiro, Brazil. medRxiv 10.1101/2020.12.23.20248598.

Walls, A.C., Park, Y.J., Tortorici, M.A., Wall, A., McGuire, A.T., and Veesler, D. (2020). Structure, Function, and Antigenicity of the SARS-CoV-2 Spike Glycoprotein. Cell 181, 281–292 e286.

Walls, A.C., Tortorici, M.A., Bosch, B.J., Frenz, B., Rottier, P.J.M., DiMaio, F., Rey, F.A., and Veesler, D. (2016). Cryo-electron microscopy structure of a coronavirus spike glycoprotein trimer. Nature 531, 114–117.

Wang, C., van Haperen, R., Gutiérrez-Álvarez, J., Li, W., Okba, N.M.A., Albulescu, I., Widjaja, I., van Dieren, B., Fernandez-Delgado, R., Sola, I., et al. (2021). A conserved immunogenic and vulnerable site on the coronavirus spike protein delineated by cross-reactive monoclonal antibodies. Nature Communications 12.

Wang, H., Gristick, H.B., Scharf, L., West, A.P., Galimidi, R.P., Seaman, M.S., Freund, N.T., Nussenzweig, M.C., and Bjorkman, P.J. (2017). Asymmetric recognition of HIV-1 Envelope trimer by V1V2 loop-targeting antibodies. Elife 6.

Wang, N., Li, S.-Y., Yang, X.-L., Huang, H.-M., Zhang, Y.-J., Guo, H., Luo, C.-M., Miller, M., Zhu, G., Chmura, A.A., et al. (2018). Serological Evidence of Bat SARS-Related Coronavirus Infection in Humans, China. Virologica Sinica 33, 104–107.

Wec, A.Z., Wrapp, D., Herbert, A.S., Maurer, D.P., Haslwanter, D., Sakharkar, M., Jangra, R.K., Dieterle, M.E., Lilov, A., Huang, D., et al. (2020). Broad neutralization of SARS-related viruses by human monoclonal antibodies. Science 369, 731–736.

Weisblum, Y., Schmidt, F., Zhang, F., DaSilva, J., Poston, D., Lorenzi, J.C.C., Muecksch, F., Rutkowska, M., Hoffmann, H.-H., Michailidis, E., et al. (2020). Escape from neutralizing antibodies by SARS-CoV-2 spike protein variants. eLife 9.

West, A.P., Barnes, C.O., Yang, Z., and Bjorkman, P.J. (2021). SARS-CoV-2 lineage B.1.526 emerging in the New York region detected by software utility created to query the spike mutational landscape. bioRxiv 10.1101/2021.02.14.431043;.

West, A.P., Jr., Scharf, L., Horwitz, J., Klein, F., Nussenzweig, M.C., and Bjorkman, P.J. (2013). Computational analysis of anti-HIV-1 antibody neutralization panel data to identify potential functional epitope residues. Proc Natl Acad Sci U S A 110, 10598–10603.

Winn, M.D., Ballard, C.C., Cowtan, K.D., Dodson, E.J., Emsley, P., Evans, P.R., Keegan, R.M., Krissinel, E.B., Leslie, A.G., McCoy, A., et al. (2011). Overview of the CCP4 suite and current developments. Acta Crystallogr D Biol Crystallogr 67, 235–242.

Wrapp, D., Wang, N., Corbett, K.S., Goldsmith, J.A., Hsieh, C.L., Abiona, O., Graham, B.S., and McLellan, J.S. (2020). Cryo-EM structure of the 2019-nCoV spike in the prefusion conformation. Science 367, 1260–1263.

Yan, R., Zhang, Y., Li, Y., Xia, L., Guo, Y., and Zhou, Q. (2020). Structural basis for the recognition of SARS-CoV-2 by full-length human ACE2. Science 367, 1444–1448.

Yuan, M., Liu, H., Wu, N.C., Lee, C.-C.D., Zhu, X., Zhao, F., Huang, D., Yu, W., Hua, Y., Tien, H., et al. (2020a). Structural basis of a shared antibody response to SARS-CoV-2. Science 10.1126/science.abd2321, eabd2321.

Yuan, M., Wu, N.C., Zhu, X., Lee, C.-C.D., So, R.T.Y., Lv, H., Mok, C.K.P., and Wilson, I.A. (2020b). A highly conserved cryptic epitope in the receptor binding domains of SARS-CoV-2 and SARS-CoV. Science 368, 630–633.

Yuan, Y., Cao, D., Zhang, Y., Ma, J., Qi, J., Wang, Q., Lu, G., Wu, Y., Yan, J., Shi, Y., et al. (2017). Cryo-EM structures of MERS-CoV and SARS-CoV spike glycoproteins reveal the dynamic receptor binding domains. Nat Commun 8, 15092.

Zhang, W., Davis, B.D., Chen, S.S., Sincuir Martinez, J.M., Plummer, J.T., and Vail, E. (2021). Emergence of a Novel SARS-CoV-2 Variant in Southern California. Jama 10.1001/jama.2021.1612.

Zhou, D., Duyvesteyn, H.M.E., Chen, C.-P., Huang, C.-G., Chen, T.-H., Shih, S.-R., Lin, Y.-C., Cheng, C.-Y., Cheng, S.-H., Huang, Y.-C., et al. (2020a). Structural basis for the neutralization of SARS-CoV-2 by an antibody from a convalescent patient. Nature Structural & Molecular Biology 27, 950–958.

Zhou, H., Ji, J., Chen, X., Bi, Y., Li, J., Hu, T., Song, H., Chen, Y., Cui, M., Zhang, Y., et al. (2021). Identification of novel bat coronaviruses sheds light on the evolutionary origins of SARS-CoV-2 and related viruses. bioRxiv 10.1101/2021.03.08.434390.

Zhou, P., Yang, X.L., Wang, X.G., Hu, B., Zhang, L., Zhang, W., Si, H.R., Zhu, Y., Li, B., Huang, C.L., et al. (2020b). A pneumonia outbreak associated with a new coronavirus of probable bat origin. Nature 579, 270–273.

Zivanov, J., Nakane, T., Forsberg, B.O., Kimanius, D., Hagen, W.J., Lindahl, E., and Scheres, S.H. (2018). New tools for automated high-resolution cryo-EM structure determination in RELION-3. Elife 7.

Zost, S.J., Gilchuk, P., Case, J.B., Binshtein, E., Chen, R.E., Nkolola, J.P., Schafer, A., Reidy, J.X., Trivette, A., Nargi, R.S., et al. (2020a). Potently neutralizing and protective human antibodies against SARS-CoV-2. Nature 584, 443–449.

Zost, S.J., Gilchuk, P., Chen, R.E., Case, J.B., Reidy, J.X., Trivette, A., Nargi, R.S., Sutton, R.E., Suryadevara, N., Chen, E.C., et al. (2020b). Rapid isolation and profiling of a diverse panel of human monoclonal antibodies targeting the SARS-CoV-2 spike protein. Nat Med 10.1038/s41591-020-0998-x.

